# Intraperitoneal programming of tailored CAR macrophages via mRNA-LNP to boost cancer immunotherapy

**DOI:** 10.1101/2024.07.30.605730

**Authors:** Kedan Gu, Ting Liang, Luting Hu, Yifan Zhao, Weiyang Ying, Mengke Zhang, Yashuang Chen, Benmeng Liang, Xinrui Lin, Yanqi Zhang, Hongyu Wu, Meng Wang, Yuping Zhu, Wenxi Wang, Yu Zhang, Chao Zuo, Zhen Du, Penghui Zhang, Jia Song, Xiangsheng Liu, Sitao Xie, Weihong Tan

## Abstract

Therapeutic strategies for peritoneal metastasis in solid tumors are urgently needed in the clinic. Programming chimeric antigen receptor macrophages (CAR-Ms) *in situ* offers opportunities for an unmet demand. However, potential intracellular domains (ICDs) for CAR design and their antitumor mechanisms for macrophage empowerment remain to be explored systematically. By developing a targeted mRNA-LNP delivery system for macrophages, we have investigated 36 CAR combinations to determine the impact of CAR-Ms on immune regulation *in vitro* and *in vivo*. In two solid tumor mouse models, intraperitoneal programming of CAR-Ms was shown to elicit robust adaptive immune activation and significantly synergize with PD-1/L1 therapy. Single-cell RNA sequencing (scRNA-seq) analysis revealed that CAR-Ms could reshape the immunosuppressive tumor microenvironment (TME) and boost the TCF1^+^PD-1^+^ progenitor-exhausted CD8^+^ T cells (Tpex) population. Meanwhile, we found that tailored CAR-M with CD3ζ/TLR4 ICDs could favorably maintain proinflammatory phenotype and simultaneously upregulate MHC I and PD-L1 expression by perturbing NF-κB pathways. Moreover, the synergism between macrophage PD-L1 knockdown and CAR-M therapy highlighted the need to block the PD-1/L1 axis in antigen cross-presentation. In short, we developed an mRNA-LNP delivery system for intraperitoneal programming of tailored CAR-Ms *in vivo* and broadened understanding of both regulatory and feedback mechanisms for CAR-M therapies against solid tumors.

## INTRODUCTION

Peritoneal metastasis in solid tumors poses significant clinical challenges in oncology^1^. A typical combination of cytoreductive surgery (CRS) and hyperthermic intraperitoneal chemotherapy (HIPEC) provides benefits to a minority of patients with minimal tumor burdens^1,2^. At the same time, it is not considered a clinical option for the majority of patients with advanced peritoneal tumors^3,4^. Immunotherapy represents a promising frontier for addressing peritoneal metastasis^5^. However, peritoneal tumors often develop evasive mechanisms against the immune system, resulting in disease progression and unfavorable outcomes^6,7^. Hence, there is a clinically urgent and unmet need to explore alternative immunotherapies for the majority of peritoneal metastasis patients.

Meanwhile, peritoneal ascites from peritoneal metastasis patients harbor a substantial population of immune cells, with macrophages constituting approximately 45%^8^. As for the tumor microenvironment (TME), tumor-associated macrophages (TAMs) are roughly categorized into proinflammatory M1-like or pro-tumoral M2-like up to 50% of total cells^9,10^. High levels of M2-like TAM infiltration are often associated with poor prognosis and resistance to immunotherapy in clinical trials^11,12^. Therefore, approaches to altering the phenotype or function of macrophages to enhance immune responses against solid tumors are under investigation^13–20^.

In particular, CAR macrophages (CAR-Ms) exhibit a pronounced phagocytic activity toward target cells and can easily penetrate solid tumors, demonstrating decreased tumor burden and prolonged overall survival in preclinical solid tumor models^21–27^. However, the elaborate and costly manufacturing processes similar to those FDA-approved CAR-T^21,25,28^ , when coupled with the tumorigenic risk^29,30^ of viral vector-involved cell engineering, restrict the accessibility of CAR-Ms to patients who would otherwise benefit from the broader clinical applications. In response, researchers began trying to generate CAR-Ms *in vivo* to treat solid tumors directly by constructing non-viral nanocarriers^31^. Furthermore, current research on the architecture of the intracellular domain (ICD) of macrophages primarily focuses on CD3ζ signal transduction. Yet, exclusive reliance on CD3ζ is far from fully exploiting the multifunctional characteristics of macrophages for tumor immunotherapies. One could speculate on the utility of T cell-based ICDs, but these may not effectively function in macrophages for solid tumors. Therefore, systematically investigating the rational design, combination, and biological effects of signaling pathways implicated in macrophages through the CAR modality is in order.

Therefore, in this work, we explored the biological effects of 36 CAR combinations containing diverse ICDs (Phagocytosis: CD3ζ and Dectin1; proinflammatory: CD40 and TLR4; and possible effector: CD46 and CFS2R) on macrophages *in vitro* and *in vivo*. By developing a targeted mRNA-LNP delivery system for macrophages, we achieved highly efficient *in vivo* construction of CAR- M required for cancer immunotherapy. In two syngeneic solid tumor mouse models, intraperitoneal programming of tailored CAR-Ms elicited robust adaptive immune system activation and significantly synergizes with standard-of-care PD-1/L1 immune checkpoint blockade (ICB) therapy in resistant models. Further analysis by comprehensive single-cell RNA sequencing (scRNA-seq) demonstrated that *in vivo* programming of CAR-Ms with CD3ζ/TLR4 ICDs could significantly promote the transition of macrophages from M2 to M1 proinflammatory phenotype, accompanied by perturbations of the NF-κB pathway to upregulate PD-L1 and major histocompatibility complex class I (MHC I). Meanwhile, we found that CAR-Ms reshaped the immunosuppressive tumor microenvironment, thereby boosting the population of TCF1^+^PD-1^+^ progenitor exhausted CD8 T cells (Tpex). Moreover, the synergistic effect between siRNA-mediated knockdown of PD-L1 and CAR-Ms highlighted the critical role of PD-L1 expression on macrophages in the antigen cross-presentation process.

## RESULTS

### Functional design and optimization of CARs with variable signal transduction ICDs for CAR-M

Orchestrated through an array of natural receptors, macrophages sense and respond to external cues, executing multifaceted roles that encompass phagocytosis, immunomodulation and antigen presentation. Noteworthy among these receptors, the recognition of β-glucans by the C-type lectin receptor Dectin1 underscores the pivotal role of macrophages in fungal defense mechanisms^32,33^. By engaging with its ligand CD40L, CD40 receptor facilitates transition towards the proinflammatory M1 phenotype^34,35^, thereby asserting immune homeostasis. Additionally, recognition of lipopolysaccharides (LPS) by Toll-like receptors TLR2 and TLR4 triggers intracellular signaling cascades^36^ that culminate in activating proinflammatory pathways. CD46 is a well-known viral magnet, and CSF2R is common to IL-3, IL-5 and GM-CSF receptors. However, whether these natural receptor ICDs could be designed to functionalize CAR-M in a manner similar to CD3ζ remains to be elucidated. Thus, we have meticulously developed a repertoire of CARs integrating a spectrum of signal transduction ICDs (CD40, CD46, CSF2R, Dectin1, and TLR4) tailored to macrophages with the aim of manipulating macrophage functions via defined antigenic target stimulation (Fig. 1a and Supplementary Table 1).

**Fig. 1.**
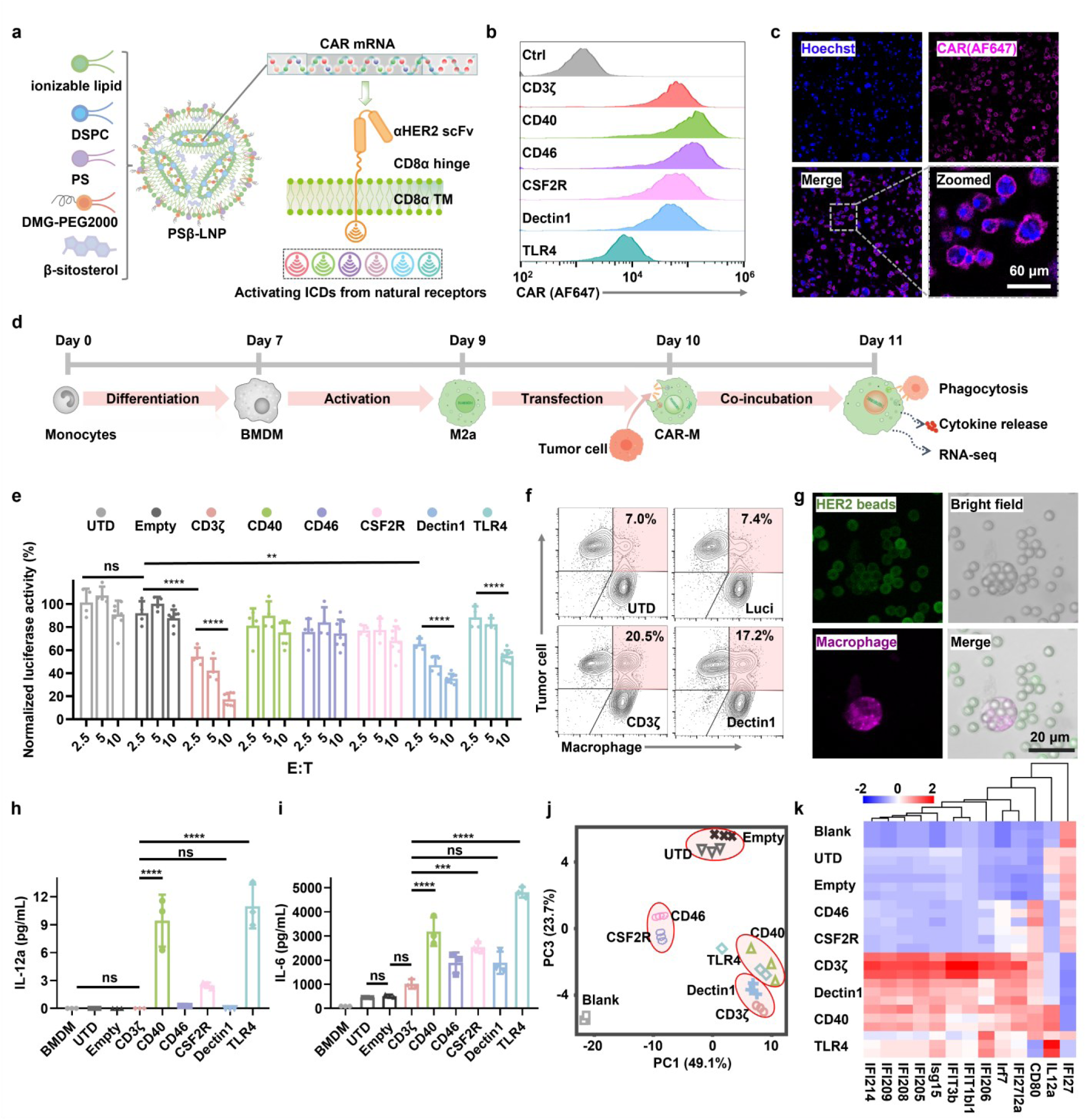
| Design and functionality of CARs with variable ICDs for CAR-M. a, Schematic illustration of *in vitro* preparation of CAR-M containing pending CARs via mRNA PSβ-LNP. PSβ, phosphatidylserine and β-sitosterol. Flow cytometry analysis **(b)** and confocal images **(c)** of CARs expression on bone marrow-derived macrophages (BMDM) 24 h after mRNA PSβ-LNP transfection. CAR and nuclei were counterstained with HER2-AF647 (magenta) and Hoechst33258 (blue), respectively. Scale bar, 60 μm for the zoomed image. **d,** Schematic procedures for functionality verification of CAR-M with pending CARs. **e,** Normalized phagocytosis of targeted luciferase-reported tumor cells by anti-HER2 CAR-Ms with pending ICDs in a series of E:T ratios at 2.5, 5 and 10. Data are shown in means ± SD (n ≥ 4). Statistical significance was calculated using two-way ANOVA. **f,** Flow cytometry analysis of adhesion of controls or CAR-Ms and tumor cells 4 h after coincubation. **g,** Confocal images of phagocytosis of HER2-beads by anti-HER2 CAR (CD3ζ) expressing THP-1 macrophage. HER2 beads and CAR-expressing THP-1 were stained with FITC (green) and DiD (magenta), respectively. Scale bar, 20 μm. Detected cytokines IL-12 **(h)** and IL-6 **(i)** in medium supernatant of controls or CAR- Ms and tumor cells 24 h after coincubation. Statistical significance was calculated using one-way ANOVA. Gene expression PCA **(j)** and heatmap **(k)** from bulk RNA-seq (n = 3) clustering from Blank, UTD, Empty, and CAR-Ms with ICDs (CD3ζ, CD40, CD46, CSF2R, Dectin1, and TLR4). For all panels, ns = no significance, \**P* < 0.05, \*\**P* < 0.01, \*\*\**P* < 0.001, \*\*\*\**P* < 0.0001.

To avoid immune responses to foreign nucleic acids caused by intracellular immune sensors in macrophages, all designed mRNAs were modified with N1-methyl-pseudouridine to lower immunogenicity^37^. Phosphatidylserine, as the macrophage-specific uptake unit^38^, and β-sitosterol to improve mRNA transfection^39^ were incorporated in the existing mRNA-based lipid nanoparticles (LNP) delivery system to form phosphatidylserine/β-sitosterol LNP (termed “PSβ- LNP”) (Fig. 1a), which could perform targeted delivery of mRNA to macrophages with high efficiency (Supplementary Fig. 1a). High transfection efficiency was achieved *in vitro* among various macrophage cell types, including THP-1, RAW264.7 and bone marrow-derived macrophages (BMDMs) for both CAR-mRNA (Fig. 1b) and eGFP-mRNA (Supplementary Fig. 1b). Confocal imaging indicated that the mRNA-encoded CARs could be localized on the cell membrane (Fig. 1c). PSβ-LNP was characterized as having an average diameter of 164 ± 0.7 nm, a polydispersity index (PDI) of less than 0.1, and a pKa of around 6.72, demonstrating a controllable particle size distribution and outstanding endosomal escape capabilities at the cellular level (Supplementary Fig. 1c-1f).

To ascertain the functionality potentially mediated by the designed CARs, CAR-Ms were generated from transfected BMDMs using corresponding CAR mRNA encapsulated within the PSβ-LNP. Subsequently, macrophages and luciferase-expressing tumor cells were cocultured and analyzed as depicted in Fig. 1d. It should be noted that Dectin1 CAR-M eradicated tumor cells in an E:T (effector cells: target cells) ratio-dependent manner similar to that of CD3ζ CAR-M (Fig. 1e), further noting that its phagocytic activity was not dependent on the amount of CAR-encoding mRNA (Supplementary Fig. 2a). Additional imaging analysis showed that the pHrodo^TM^ Red-labeled tumor cells emitted bright red fluorescence in the CD3ζ or Dectin1 CAR-M group, suggesting that engulfed tumor debris was degraded in acidic phagolysosome lumen (Supplementary Fig. 2b). Flow cytometry data indicated a more vital interaction between CD3ζ or Dectin1 CAR-Ms and tumor cells by the proportion of calculated changes of 20.5% or 17.4%, respectively, compared to the untreated (UTD) group of 7.0% (Fig. 1f). Antigen-specific beads engulfment assay further demonstrated successful CAR-M phagocytosis (Fig. 1g and Supplementary Fig. 2c-2e). Furthermore, we observed a marked elevation of proinflammatory cytokines, including IL-6, IL-12 and TNF-α, in the supernatant of the CD40 or TLR4 CAR-M treated group (Fig. 1h, 1i and Supplementary Fig. 2f). Next, CAR-Ms were further analyzed to explore the relationship between CAR effects and gene changes. RNA-seq principal component analysis (PCA) and featured interferon-associated genes heatmap revealed that CD3ζ showed a gene signature similar to that of Dectin1, whereas CD40 was similar to TLR4 (Fig. 1j, 1k and Supplementary Fig. 3a). Heightened expression of the IL-12 gene was only detected in the CD40 and TLR4 groups in alignment with the findings from cytokine analysis in the supernatant (Fig. 1h and 1k). The stimulation induced by empty LNP, deemed negligible, was similar to that of the UTD group in the coculture system. Neither the CD46 nor CSF2R CAR design appeared to mediate much discernible functionality since no significant difference was observed compared to that of either UTD or Empty groups, especially in featured genes (Fig. 1j and 1k). Gene ontology (GO) analysis suggested that four functional CARs for CAR-M mediated activation of innate immune response (Supplementary Fig. 3). Taken together, CD3ζ or Dectin1 CARs primarily mediated macrophage with enhanced phagocytosis functionalities, while CD40 or TLR4 CARs induced more marked proinflammatory regulation.

To improve the antitumor performance of CAR-M, we further optimized the ICD and extracellular domain (ECD) of CARs. Considering the temporal limitation of mRNA-encoding CAR and the potential degradation or loss of CAR structure after target antigen recognition, we introduced a lysine-to-arginine (K-R) mutation in the ICD (Fig. 2a and Supplementary Table 1), a strategy inspired by CAR-T to prolong the circulation time of CAR^40^. CAR^K-R^ and CAR^K^ exhibited similar metabolic profiles in the presence of target tumor cells. Despite the consistently higher positivity rate of CAR^K-R^ at each time point of detection, the difference compared to the CAR^K^ was statistically marginal (Fig. 2b and 2c). As for ECD optimization, given the revealed coordinative transcriptional regulation and expression of CD47 and HER2^41,42^, as well as the pivotal role of the CD47-SIRPα axis in the inhibition of macrophage phagocytosis, we engineered mutated SIRPα^43^ as a high-affinity CD47 antigen recognition domain of the CAR construct to disrupt CD47-SIRPα interaction in order to liberate macrophage cytotoxicity (Fig. 2a). With an E:T ratio of 10:1, the αCD47 CAR markedly augmented CAR-M phagocytosis to near-eradication of all tumor cells, whereas about 25% of tumor cells remained alive in the αHER2 CAR-M group (Fig. 2d and Supplementary Fig. 4a). With CD47- and HER2-positive or -negative tumor cells as target, it is confirmed that phagocytic activation could potentially be triggered by stimulation through either CD47 or HER2 (Fig. 2e-2g and Supplementary Fig. 4b). IncuCyte-based assays demonstrated the continuous and specific phagocytic process up to 72 hours (Fig. 2h, 2i and Supplementary movie).

**Fig. 2.**
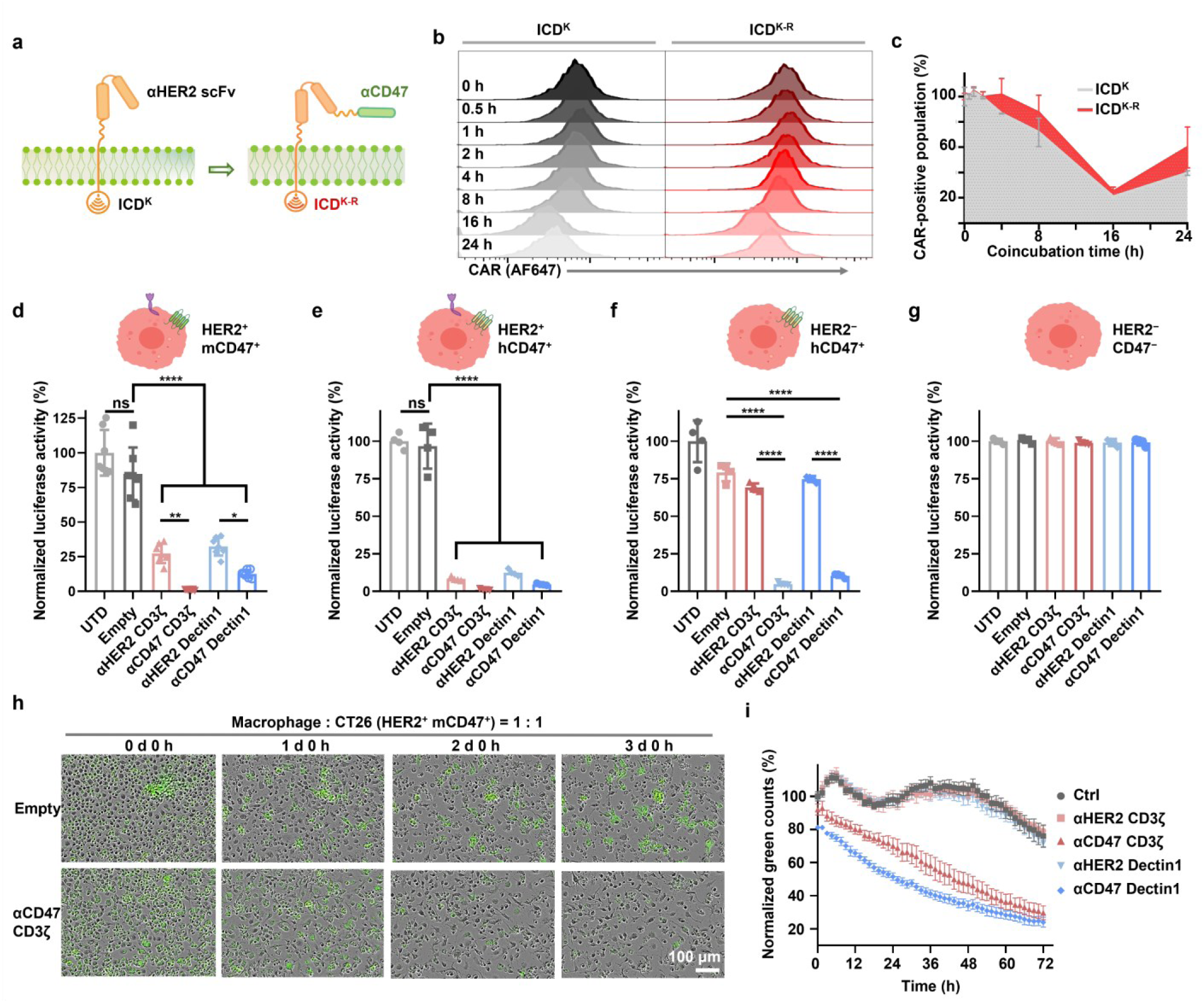
| Mutation of CAR ICD for enhancing persistence and CD47 recognition of CAR ECD for enhancing phagocytosis. a, Schematic illustration of CAR optimization. **b,** Flow cytometry analysis of CAR expression of ICD^K^ and ICD^K-R^. **c,** Positive CAR-Ms with ICD^K^ or ICD^K-R^ during continued target cell stimulation. Normalized phagocytosis of luciferase-reported target tumor cells (**d,** CT26-huERBB2^+^-mCD47^+^; **e,** SKOV3-huERBB2^+^-huCD47^+^; and **f,** HS578T-huERBB2^−^-huCD47^+^) by CAR-Ms. E:T = 10:1. **g,** Normalized phagocytosis of luciferase-reported non-target tumor cells (MDA-MB-231-huERBB2^−^-CD47^−^) by CAR-Ms. E:T = 10:1. **h,** Incucyte-based phagocytosis pictures of CT26-huERBB2^+^-mCD47^+^-eGFP^+^ by αCD47(CD3ζ) CAR-M or empty control within three days. E:T = 1:1. **i,** Incucyte-based phagocytosis assay of HS578T-huERBB2^−^-huCD47^+^-eGFP^+^ by CAR-Ms or control within three days. For all panels, statistical significance was calculated using one-way ANOVA. For all panels, ns = no significance, \**P* < 0.05, \*\**P* < 0.01, \*\*\**P* < 0.001, \*\*\*\**P* < 0.0001.

### Intraperitoneal programming of CAR-M via mRNA PSβ-LNP

It was found that peritoneal injection of CAR-Ms mainly enriched in tumor tissue^44^. Concurrently, considering the various barriers and complexities of metabolic distribution associated with intravenous infusion^45,46^, we speculate that it should be a more feasible way to administer mRNA PSβ-LNP for CAR-M therapy intraperitoneally. Initially, we validated the nucleic acid delivery capability of GFP-mRNA PSβ-LNP towards macrophages in a colon cancer CT26 syngeneic mouse model with peritoneal metastasis according to the protocol outlined in Fig. 3a. In malignant ascites, more than 25% GFP and F4/80 double-positive cells were detected in the PSβ-LNP group, indicating that approximately 60% F4/80-positive cells could be successfully transfected within 18 hours in a single administration, significantly surpassing the efficiency observed with traditional LNPs (Fig. 3b and 3c). Comparable results could also be observed in the pancreatic cancer PAN02 syngeneic mouse model (Supplementary Fig. 5a). Additionally, we observed that the peritoneal fluid in the empty LNP treatment group or the mRNA-LNP treatment group contained a higher total cell count compared to the control group (40∼65 million vs. 20 million) (Supplementary Fig. 5b) with a corresponding increase in the proportion of macrophages from 37.8% to 45.6% or 54.6% (Fig. 3b). For malignancy itself, approximately 23% of intratumoral F4/80-positive cells were transfected. In contrast, three-quarters of all transfected cells were F4/80-positive (Fig. 3d), suggesting excellent selectivity and transfection ability of PSβ-LNP towards macrophages. Regarding the spatial distribution of transfected cells, further immunofluorescent (IF) sections revealed that widespread eGFP expression primarily appeared in macrophages within the tumor, ranging from the periphery to the interior (Fig. 3e). Considering the gap drop between mRNA-LNP uptake and protein of interest expression, we additionally tested the organ distribution of PSβ-LNP intraperitoneally. DIR-labeled PSβ-LNP predominantly accumulated at the tumor lumps following intraperitoneal injection with little dispersion out of the peritoneal cavity and distribution in the liver (Fig. 3f), revealing the excellent selectivity of PSβ- LNP at the organ level. Similar results were also observed in the ovarian cancer SKOV3 xenograft mouse model (Supplementary Fig. 5c and 5d).

**Fig. 3.**
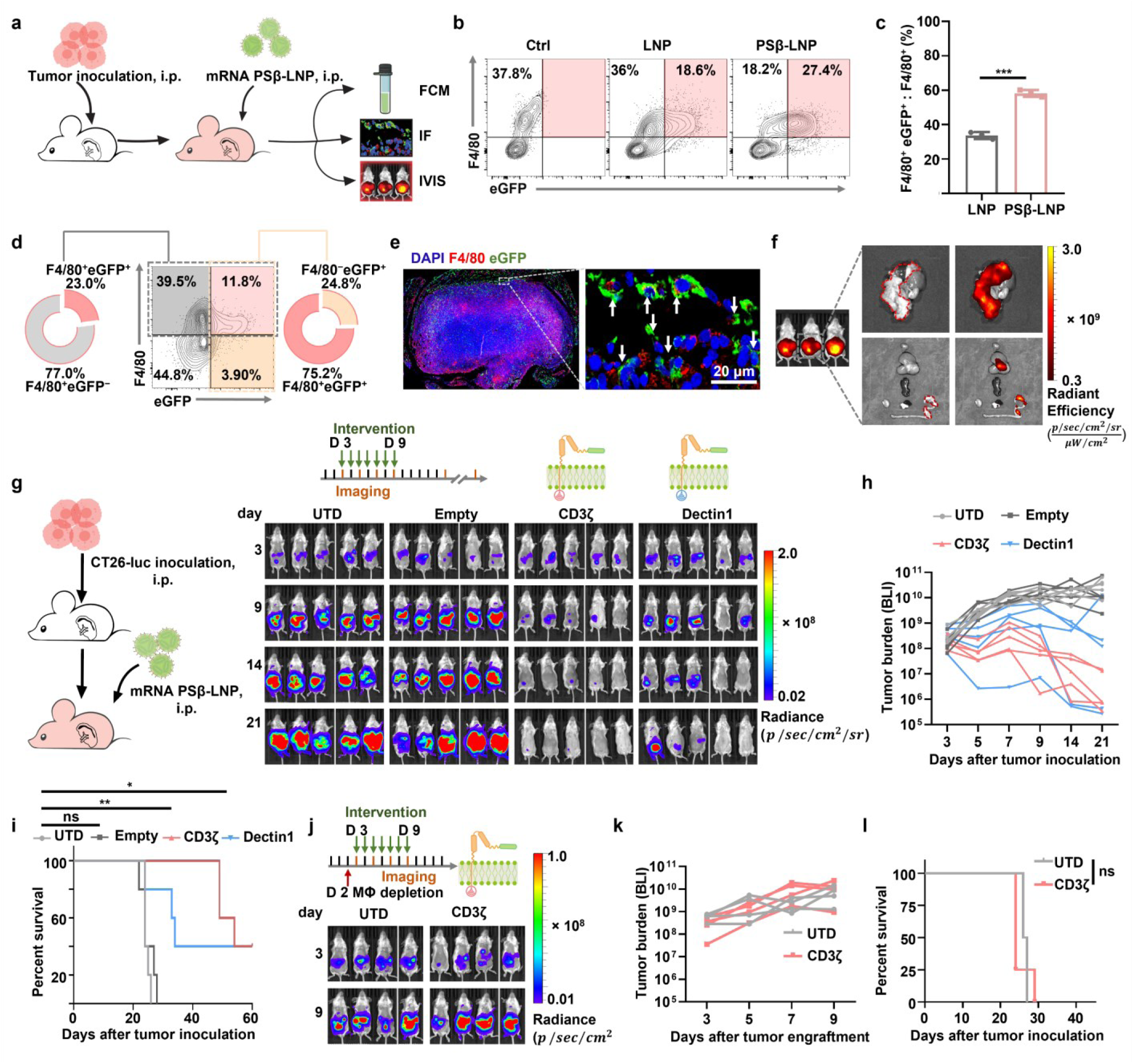
| *In situ* programming of CAR-M via intraperitoneal administration of CAR-mRNA PSβ-LNP for tumor suppression. a, Schematic experimental validation of PSβ-LNP targeting peritoneal macrophages. **b,** Flow cytometry analysis of the proportion of eGFP-expressing ascites after infusion of eGFP-mRNA LNP. **c,** eGFP and F4/80 double-positive proportion of the F4/80-positive population in b. Statistical significance was calculated using one-way ANOVA. **d,** Flow cytometry analysis of eGFP-positive intratumoral proportion after infusion of eGFP-mRNA LNP. **e,** Immunofluorescence slice (F4/80, red; eGFP, green; nuclei, blue) of tumor tissue after eGFP-mRNA PSβ-LNP infusion. Scale bar, 20 μm. **f,** DIR signal distribution after intraperitoneal administration of DIR-labeled PSβ-LNP. **g,** Schematic experimental design of CAR-mRNA PSβ-LNP programming of CAR-M *in situ* and *in vivo* bioluminescent images. Quantified signal intensity **(h)** of CT26-luc syngeneic murine model. **i,** Kaplan-Meier survival curves of mice after each treatment (n = 5). Data were analyzed using the log-rank (Mantel- Cox) test (n = 5). *In vivo* bioluminescent images **(j)** and quantified signal intensity **(k)** of CT26- luc syngeneic murine model with macrophage depletion. **l,** Kaplan-Meier survival curves of mice after each treatment (n = 4) with macrophage depletion. Data were analyzed using the log-rank (Mantel-Cox) test (n = 4). For all panels, ns = no significance, \**P* < 0.05, \*\**P* < 0.01, \*\*\**P* < 0.001, \*\*\*\**P* < 0.0001.

Encouraged by the above *in situ* transfection assay results, we further monitored the antitumor capacity of CAR-mRNA encapsulated within PSβ-LNP in the CT26-luc syngeneic mouse model (Fig. 3g). Results revealed potent tumor inhibitory effects for both CD3ζ CAR-M and Dectin1 CAR-M, significantly prolonging the survival of tumor-bearing mice (Fig. 3h and 3i). It should be noted that CD3ζ CAR demonstrated superior therapeutic efficacy compared to Dectin1 CAR, while no observable therapeutic effects were achieved in the Empty LNP group. During the early stages of treatment, the CAR-M group experienced a brief period of weight loss, followed by sustained growth (Supplementary Fig. 6a), indicating a minor and tolerable toxicity from mRNA-LNP. In the later stages of observation, control mice exhibited significant abdominal distension and rough fur. At the same time, the CAR-M treatment group did not display these symptoms (Supplementary Fig. 6b). Prior depletion of macrophages in mice using clodronate liposomes before intervention abolished the therapeutic benefits of subsequent CAR-mRNA PSβ- LNP administration (Fig. 3j-3l), underscoring the essential role of macrophages in the programming of CAR-M using CAR-mRNA PSβ-LNP *in vivo*. In summary, we comprehensively validated the *in vivo* macrophage-targeting capability, efficient mRNA delivery of PSβ-LNP, and the preliminary antitumor efficacy of CAR-mRNA LNP.

### CAR-Ms could activate the adaptive immune system and sensitize αPD-1 therapy

Considering this pivotal role of macrophages in the intricate innate and adaptive immune response network, we conducted further investigations to evaluate the influence of *in situ* generation of CAR-M on the immune system. To investigate the potential of CAR-M intervention in establishing sustained adaptive immune protection, we re-challenged mice, which had previously achieved complete tumor eradication following CAR-M treatment, by administering one million tumor cells subcutaneously (Fig. 4a). Over a prolonged observation period, these tumors were once again wholly inhibited (Fig. 4a and 4b), confirming comprehensive adaptive immune protection in mice. Additionally, to elucidate the impact of CAR-M intervention on antigen-specific T cells, we harvested splenocytes from mice and stimulated them with tumor cells for an enzyme-linked immunospot assay (ELISPOT) assay. Based on ELISPOT assay, the number of tumor-specific T cells in cured mice was significantly elevated compared to naive mice (1,118 vs. 253 tumor-specific T cells detected per 1.5 × 10^5^ splenocytes) (Fig. 4c), indicating that CAR-M intervention enhances the proliferation of antigen-specific T cells. Moreover, the adoptive transfer of 10 million splenocytes from these cured mice to naive mice resulted in a pronounced inhibition of tumor growth (Fig. 4d). These experimental results provide compelling evidence for the activation of adaptive immunity by *in situ* CAR-M. However, once the tumor microenvironment (TME) was fully established, the adoptive transfer of splenocytes from cured mice demonstrated only a marginal tumor suppression (Fig. 4e), which could be partly attributed to the limited ability of tumor-specific T cells to infiltrate the tumor interior (Fig. 4f).

**Fig. 4.**
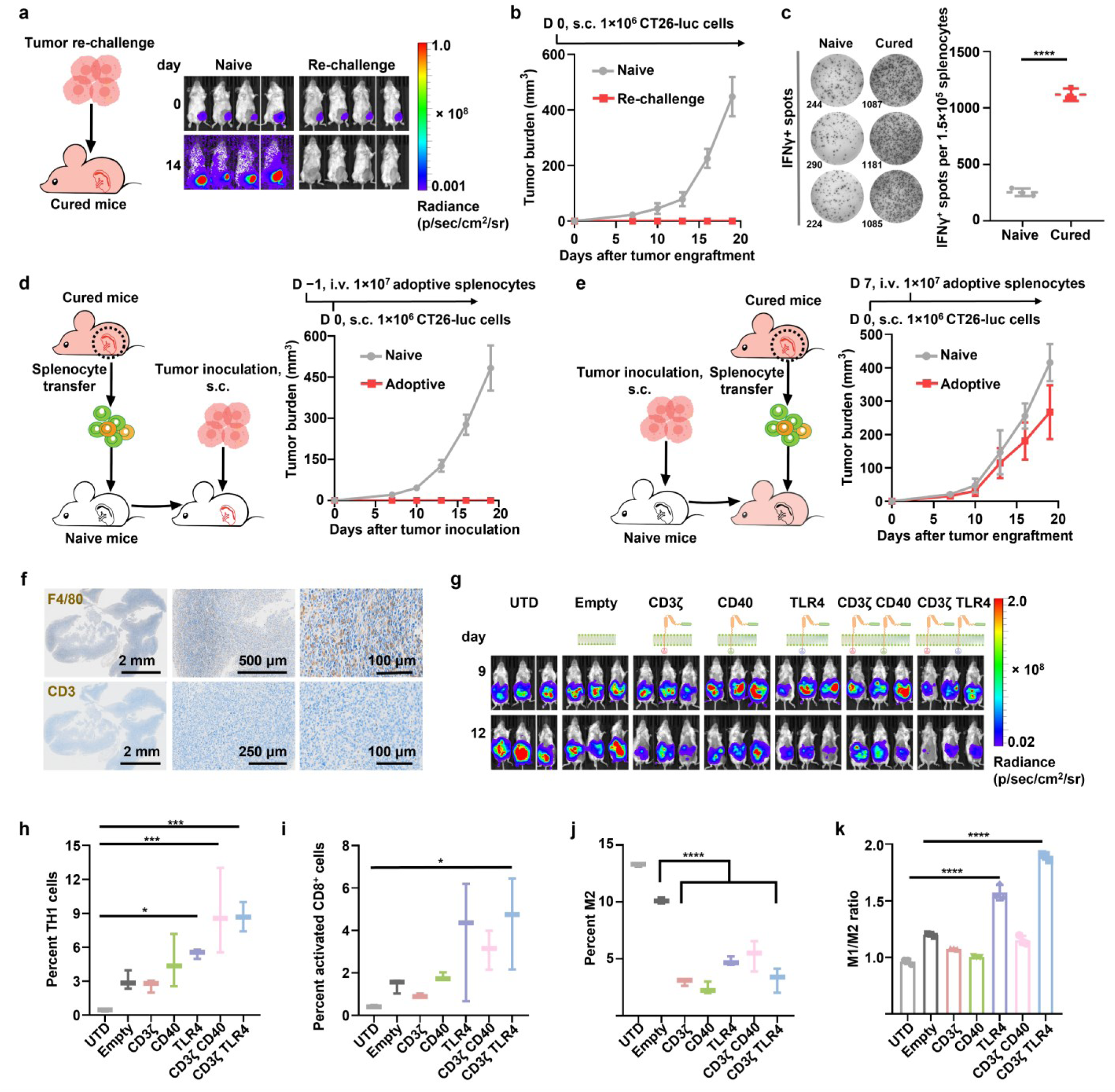
| *In situ* programming of CAR-Ms activates the adaptive immune system. *In vivo* bioluminescent images **(a)** and tumor volume **(b)** of CT26-luc syngeneic rechallenged mice or naive mice taken on the 65th day after prior CT26-luc inoculation. **c,** Representative spot pictures and quantification data in IFNγ ELISPOT assay. Statistical significance was determined with a one-tailed unpaired t-test. **d,** Tumor volume of mice with adoptive splenocytes from CT26-luc syngeneic cured mice or naive mice. Adoptive transfer of splenocytes occurred one day before tumor cell inoculation. **e,** Tumor volume of mice with adoptive splenocytes from CT26-luc syngeneic cured mice or naive mice. Adoptive transfer of splenocytes occurred 7 days after tumor cell inoculation. **f,** Immunohistochemical analysis of CT26-luc tumor tissues resected in **(e)**. Nuclei, blue; Macrophage (top, anti-F4/80, brown); T cell (down, anti-CD3, brown). Each panel shows low (left), medium (middle), and high (right) power views. **g,** *In vivo* bioluminescent images with each treatment. **h,** Percentage of TH1 cells and **(i)** active CD8^+^ T cells in ascites with each treatment. **j,** Percentage of intratumoral M2 cells and **(k)** the ratio of M1 and M2. Statistical significance for **(h-k)** was calculated using one- way ANOVA. For all panels, ns = no significance, \**P* < 0.05, \*\**P* < 0.01, \*\*\**P* < 0.001, \*\*\*\**P* < 0.0001.

Considering the proinflammatory characteristics of CD40 CAR and TLR4 CAR in macrophages, we further combined them with the phagocytic-promoting CD3ζ and evaluated their properties in an immunocompetent mouse model with a more advanced tumor. We found that CD3ζ CAR, TLR4 CAR, and their parallel constructs all exhibited more significant tumor-suppressive effects (Fig. 4g). Notably, the peritoneal fluid from mice treated with TLR4 CAR or the parallel sequences showed a marked increase in type 1 helper T cells and CD8^+^ reactive T cells (Fig. 4h and 4i), indicating that the parallel CARs effectively activated the adaptive immune system. Furthermore, the highest M1 to M2 macrophage ratio was observed in tumors treated with the parallel CARs (Fig. 4j and 4k), suggesting that the parallels promote maintenance of the M1 phenotype. A comprehensive assessment of extracellular cytokine release levels demonstrated higher levels of proinflammatory cytokine release in all parallels containing TLR4 CAR (Supplementary Fig. 7). Moreover, we noted that the parallel CD3ζ/TLR4 CARs demonstrated superior tumor phagocytic capacity *in vitro* compared to the parallel CD3ζ/CD40 CARs (Supplementary Fig. 8a and 8b). However, the underlying mechanism appears to differ from the previously observed TLR agonist-induced activation of the pBTK/CRT pathway, which resulted in calreticulin translocation to the cell membrane and enhanced the phagocytic effect of CD47 antibodies (Supplementary Fig. 8c-8e).

In light of potent activation of the adaptive immune system by tailored CARs in tandem, such as parallel CD3ζ/TLR4 CARs and parallel CD3ζ/CD40 noted above, we further explored tailored CAR-M with PD-1/PD-L1 immune checkpoint therapy. It is noteworthy that the CT26 mouse syngeneic tumor model is typically regarded as insensitive to PD-1 antibodies. However, in our hands PD-1 antibody and CAR-M treatment separately demonstrated mild antitumor growth effects in late-stage CT26 intraperitoneal tumors (Fig. 5a and 5b). Surprisingly, the combination of PD-1 antibody and CAR-M (termed as “Comb”) led to almost complete eradication during a brief dosing period and significantly improved mouse survival rates (Fig. 5a-5c). To procure sufficient tumor tissue for subsequent analysis, dosing initiation was delayed until day 12 post-

**Fig. 5.**
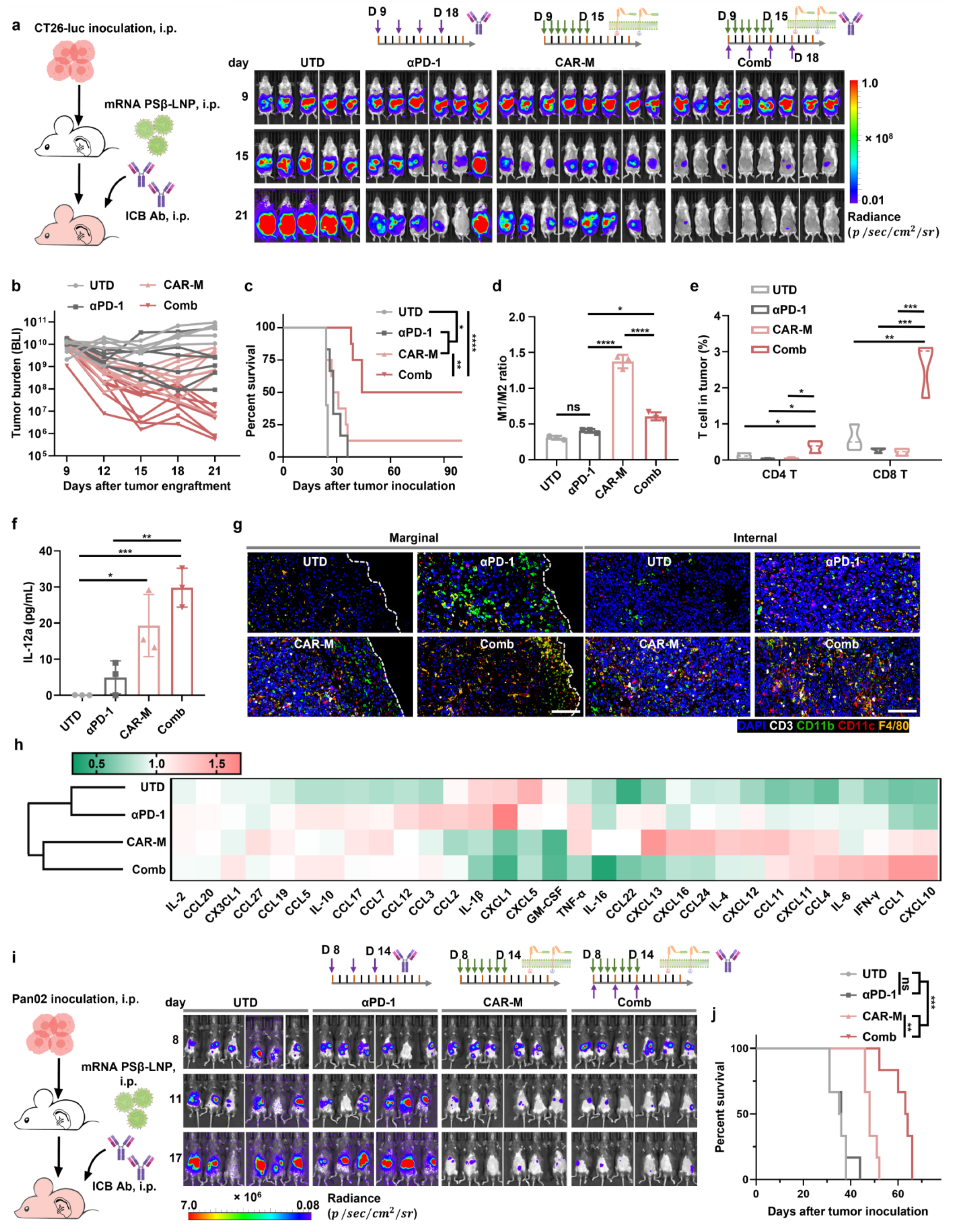
| CAR-Ms priming of αPD-1 in PD-1/PD-L1-resistant mouse model. *In vivo* bioluminescent images **(a)** and quantified signal intensity **(b)** of CT26-luc syngeneic mouse model with each treatment. **c,** Kaplan-Meier survival curves of mice after each treatment (UTD, n = 5; αPD-1, n = 6, CAR-M, n = 8; Comb, n = 8). Data were analyzed by using the log-rank (Mantel- Cox) test. **d,** The ratio of intratumoral M1 and M2. e, Percentage of intratumoral CD4^+^ or CD8^+^ T cells. Statistical significance for (**d** and **e**) was calculated using one-way ANOVA. **f,** Levels of IL- 12a in the blood across different groups. Statistical significance was calculated using one-way ANOVA. **g,** Multicolor immunofluorescence (nuclei, blue; CD3, white; CD11b, green; CD11c, red; F4/80, orange) of tumor tissue slices. Left panel is the tumor interior, and right panel is the tumor margin. **h,** Hierarchical clustering of TME cytokine concentration determined via Luminex multiplex cytokine assay under each treatment. *In vivo* bioluminescent images **(i)** and Kaplan- Meier survival curves **(j)** of the PAN02-luc syngeneic mouse model with each treatment. For all panels, ns = no significance, \**P* < 0.05, \*\**P* < 0.01, \*\*\**P* < 0.001, \*\*\*\**P* < 0.0001.

tumor inoculation. The intervention demonstrated a trend consistent with that observed previously (Supplementary Fig. 9a). Within the CAR-M treatment group, the proportion of M1 phenotype in tumor lumps was highest (Fig. 5d). In contrast, the combination treatment group showed significantly increased T-cell infiltration, especially CD8^+^ T cells (Fig. 5e and Supplementary Fig. 9b). Meanwhile, higher levels of IL-12 were detected in the blood in the CAR-M and Comb treatment groups (Fig. 5f), indicating a milieu conducive to multiple immune cells, particularly cytotoxic T cells. To comprehensively illustrate the changes in cellular composition within the TME, multicolor immunofluorescence staining was performed (Fig. 5g). In the untreated group, immune cell numbers were significantly lower, both at the core and periphery of the tumor mass, compared to other experimental groups. In the αPD-1 group, a substantial increase in myeloid cell infiltration was observed. However, the CAR-M and Comb groups exhibited diverse immune cell presence in marginal and internal tumor tissues. The wide-ranging changes in cell types are closely associated with the secretion of multiple cytokines. To demonstrate such variable cytokine secretion, we further conducted a Luminex assay for cytokine cluster analysis (Fig. 5h). In hierarchical clustering, we observed consistent trends in intertumoral cytokine release between the CAR-M and Comb with upregulation of clustered cytokines (CCL1, CCL11, CXCL10, CXCL11, and IFN-γ) compared to control, or αPD-1. Multiple studies have highlighted that IFN-γ-induced chemokines, such as CXCL10 and CXCL11, play a crucial role in eliciting chemotaxis^47–49^, especially of monocytes/macrophages and T cells within the TME. In summary, CAR-M intervention led to reshaping the cellular and cytokine landscape of the TME, fostering substantial immune activation to support T cell functionality.

Since CD47 is a pan-cancer target, we applied the same treatment in the PAN02 pancreatic cancer model. Interestingly, transient monotherapy with CAR-M alone demonstrated robust tumor eradication capabilities, while the PD-1 antibody was utterly ineffective (Fig. 5i and 5j). However, recurrence in pancreatic cancer models is common and leads to relatively low rates of complete remission. Nonetheless, we observed that the synergistic effect of CAR-M combined with PD-1 antibodies still resulted in delayed tumor recurrence and a marked extension of overall survival (Fig. 5i, 5j and Supplementary Fig. 9c). Therefore, it appeared that CAR-M could sensitize PD- 1/PD-L1 blockade therapy, making ICB a broadly applicable therapeutic option. However, the manifestation varied across different tumor types. Regarding safety, mouse weight changes were also within the normal range (Supplementary Fig. 9d and 9e). We did not detect intervention-related liver or kidney dysfunction (Supplementary Fig. 10) or organic lesions in treated mice (Supplementary Fig. 11). While we observed a reduction in red blood cell and platelet counts associated with targeting CD47, the overall red blood cell count remained within the safety baseline (Supplementary Fig. 12 and Supplementary Table 2). In short, CAR-M could activate adaptive immunity, especially parallel CD3ζ/TLR4 CARs, showing particularly strong effects. In addition, the tailored CAR-M combined with PD-1 antibodies demonstrated powerful synergistic effects and manageable toxicity in various PD-1/L1-resistance models.

### Synergism mechanism between CAR-M and ICB

To further elucidate the interaction between CAR-M and PD-1/L1 ICB therapy, single-cell RNA sequencing (scRNA-seq) was performed to analyze the changes of immune cells within the TME during the treatment of CT26 mouse syngeneic tumor model (Fig. 6a). After data filtering and quality control, we analyzed 8,467, 13,845, 13,986, and 14,933 immune cells in the UTD, αPD-1, CAR-M and Comb, respectively. Cell annotation showed shifts in immune cell proportions after various treatments with increased T cells and granulocytes in the CAR-M and combination therapy groups compared to αPD-1 monotherapy or UTD, along with a decrease in macrophage populations compared to αPD-1 treatment (Fig. 6b). Notably, macrophage cluster 1, which comprised most CAR-M and Comb, demonstrated the highest M1 scores and the lowest M2 scores (Fig. 6c-6d and Supplementary Fig. 13) characterized by the downregulation of TREM2 and the upregulation of CD80 and NOS2 (Fig. 6e-6g), and meanwhile the cluster 1 was in the minority in the αPD-1 treatment. These data align with prior findings (Fig. 4k and 5d), indicating that CAR-M effectively maintained the antitumor phenotype *in vivo*.

**Fig. 6.**
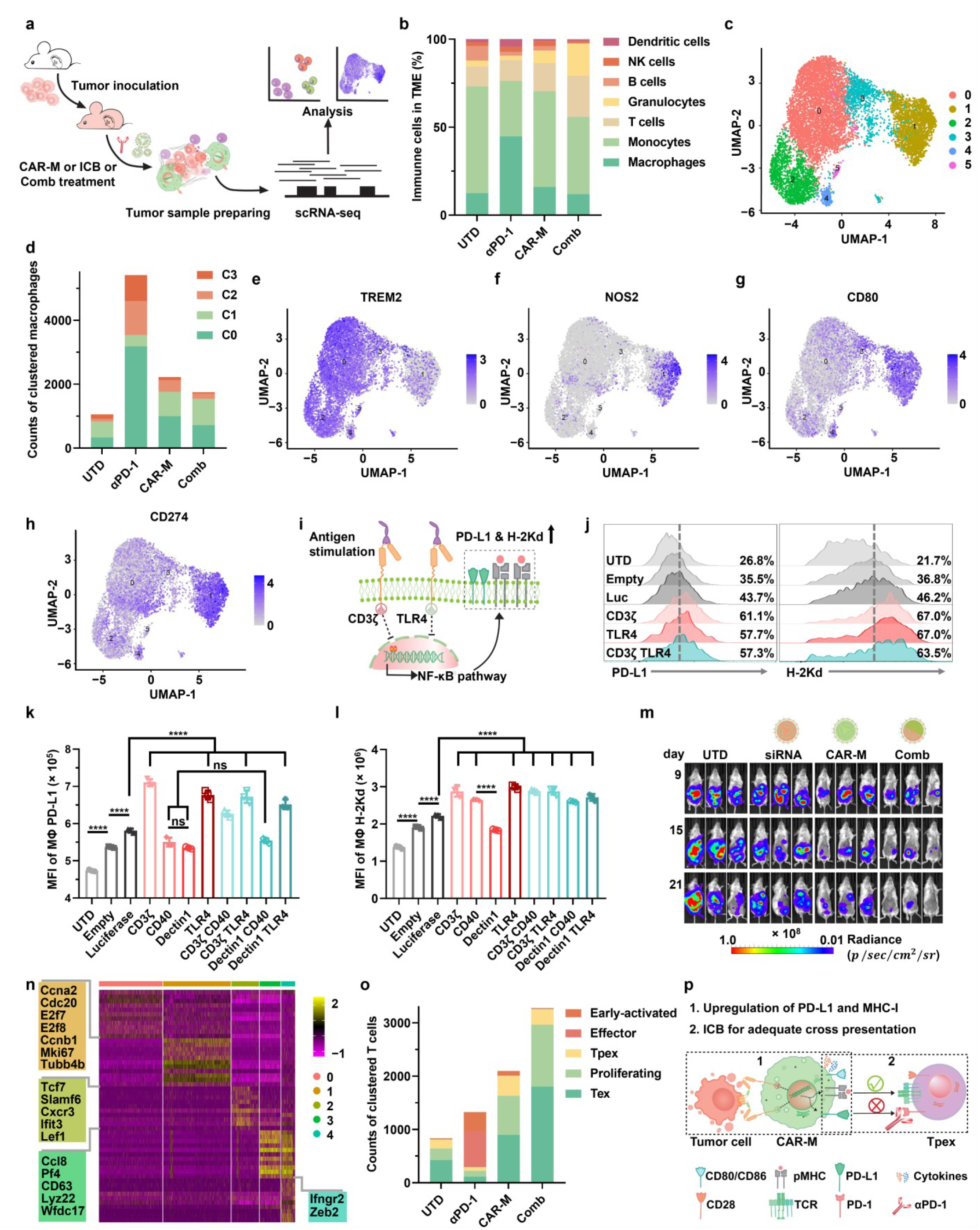
| *In situ* programming of CAR-Ms leads to M1 polarization with enhanced PD-L1 and MHC I expression. a, Schematic illustration of scRNA-seq analysis of the CT26-luc syngeneic mouse model. **b,** Proportion of tumor-infiltrating immune cells labeled via scRNA-seq. **c,** UMAP plot of scRNA-seq profiles from macrophages within the tumor. **d,** Counts of clustered macrophages in groups. UMAP plot of relative expression for the indicated gene (**e,** TREM2; **f,** NOS2; **g,** CD80; **h,** CD274). **i,** Schematic representation of CAR signal transduction leading to upregulated PD-L1 and MHC I expression. **j,** Flow cytometry analysis of macrophage PD-L1 and H-2Kd expression in groups. Mean fluorescence intensity (MFI) was calculated from flow cytometry profiles of macrophage PD-L1 **(k)** and H-2Kd **(l)**. Statistical significance was calculated using one-way ANOVA. For all panels, ns = no significance, \**P* < 0.05, \*\**P* < 0.01, \*\*\**P* < 0.001, \*\*\*\**P* < 0.0001. **m,** *In vivo* bioluminescent imaging of the CT26-luc syngeneic mouse model treated with UTD, PD-L1 siRNA, CAR-M, and combination therapy. **n,** Heatmap of featured genes of clustered tumor-infiltrating T cells. **o,** Counts of clustered T cells in groups. **p,** Schema of the mechanism of potent synergy between tailored CAR-Ms and ICB.

Indeed, studies suggested that up to 75% of tumor PD-L1 could be derived from macrophages^50,51^. We noted that macrophage cluster 1 demonstrated a significant upregulation of the PD-L1 gene (CD274) compared to other clusters (Fig. 6h), a finding consistent with our *in vitro* observations of increased PD-L1 expression in macrophages following CAR signaling (Fig. 6i-6k). Additionally, we observed a substantial upregulation of MHC I molecules and MHC I-associated gene expression in macrophages (Fig. 6i, 6j, 6l and Supplementary Fig. 14), which may contribute to the improved infiltration of CD8^+^ T cells via antigen cross-presentation within the TME. In an *in vitro* coculture system containing functional CAR-Ms and tumor cells, we found that PD-L1 expression in tumor cells was also significantly upregulated. At the same time, MHC I showed only slight changes in tumor cells (Supplementary Fig. 15). In comparison, the PD-L1 upregulation in tumor cells was nearly 10-fold greater than that in macrophages, suggesting that tumor cells demonstrate a significantly heightened negative immunoregulatory response to external interventions (Supplementary Fig. 16). Considering prior studies indicating a correlation between NF-κB pathway activation and PD-L1 and MHC I expression^52^, we reviewed previous GO (Gene Ontology) analysis data, which indicated the activation of the NF-κB pathway following CAR transduction (Supplementary Fig. 17a). Additionally, we confirmed that the CAR signaling-induced increase in PD-L1 and MHC I could be abolished by NF-κB pathway inhibitors (Supplementary Fig. 17b). While the upregulation of MHC I could enhance the efficacy of immunotherapy, its full therapeutic potential may be contingent upon concurrent PD-L1 blockade. Although disrupting the PD-L1/PD-1 axis in dendritic cells and T cells is critical, our results indicated that siRNA-mediated knockdown of substantial PD-L1 in macrophages could synergize effectively with simultaneous CAR-M therapy (Fig. 6m and Supplementary Fig. 18).

In addition to conducting a systematic and comprehensive analysis of macrophages, we examined T cells in the TME. All T cell populations within the tumor exhibited significant upregulation of exhaustion markers TOX and PDCD1 (Fig. 6n and Supplementary Fig. 19). The PD-1 antibody monotherapy group contained a higher proportion of effector T cells. In contrast, the CAR-M and Comb were enriched with terminally exhausted T cells, proliferative T cells and progenitor-exhausted T cells (Tpex) (Fig. 6o and Supplementary Fig. 19). Of particular significance was the observation that CAR-M proved advantageous in promoting the formation of Tpex clusters. Recent studies have indicated that Tpex populations are the predominant responders to PD-1 ICB^53–55^, highlighting their relevance in immunotherapy.

In summary, we elucidated the mechanisms underlying the potent synergism between tailored CAR-M and PD-1 antibodies (Fig. 6p). CAR-M, stimulated by tumor antigens, retained an M1 phenotype and upregulated MHC I and PD-L1 through the NF-κB pathway. Following the degradation of tumor cell debris by CAR-M, extensive antigen cross-presentation was performed, significantly enriching the Tpex population responsive to PD-1/PD-L1 antibodies. Moreover, the administration of PD-1 antibodies mitigated the adverse effects of PD-L1 upregulation, thereby fully liberating Tpex cells. Ultimately, CAR-M and Tpex constituted an effective cellular population against tumor cells.

## DISCUSSION

The diverse range of responses exhibited by solid tumor patients undergoing T cell-related immunotherapy highlights the necessity for exploring alternative treatment strategies. Macrophages, as critical regulators within the TME, profoundly influence therapeutic outcomes in solid tumors. Strategies for modulating the phenotype or function of macrophages to potentiate the immune microenvironment against solid tumors are currently under investigation. In peritoneal metastasis, the peritoneal cavity harbors a substantial reservoir of macrophages. Our approach, involving the intraperitoneal infusion of CAR-mRNA PSβ-LNP, facilitated the rapid and extensive generation of tumor-targeting multifunctional CAR-Ms with variable ICDs within the cavity. Consequently, upon tumor recognition, these engineered macrophages demonstrated tumor phagocytosis via CD3ζ- or Dectin1-mediated intracellular signaling, concurrently triggering the release of proinflammatory cytokines through TLR4 or CD40 signaling. This dual mechanism maintained an M1-like proinflammatory phenotype in macrophages, further activating the adaptive immune system. In two mouse syngeneic models resistant to PD-1/PD-L1 ICB, tailored CAR-Ms exhibited significant synergistic effects with PD-1 antibodies, attributed partly to TME reshaping and Tpex population expansion induced by CAR-Ms. Moreover, it was essential to emphasize that CAR signaling led to the activation of the NF-κB pathway, subsequently increasing the expression of both MHC I and PD-L1. The upregulation of MHC I on cancer cells could sensitize cancer cells to T cell-dependent killing^52^. Meanwhile, pMHC on the macrophage for presentation to T cells could serve as the primary event required for T cell activation^56^. Previous studies have emphasized the inhibitory effect of PD-L1 on dendritic cells (DCs) on T cell immunity^50,51^, while our results indicated that the PD-L1 on *in situ* prepared CAR-Ms is also crucial for adequate T cell activation, underscoring the necessity of combining CAR-Ms with PD-1 ICB therapy. In conclusion, we developed an mRNA-LNP delivery system for efficient intraperitoneal programming of tailored CAR-Ms for advanced peritoneal-disseminated late-stage tumors. *In situ* programming of CAR-Ms exhibits promising therapeutic potential, demonstrating powerful synergistic effects with traditional ICB therapy and significant promise for clinical translation.

**Scheme.**
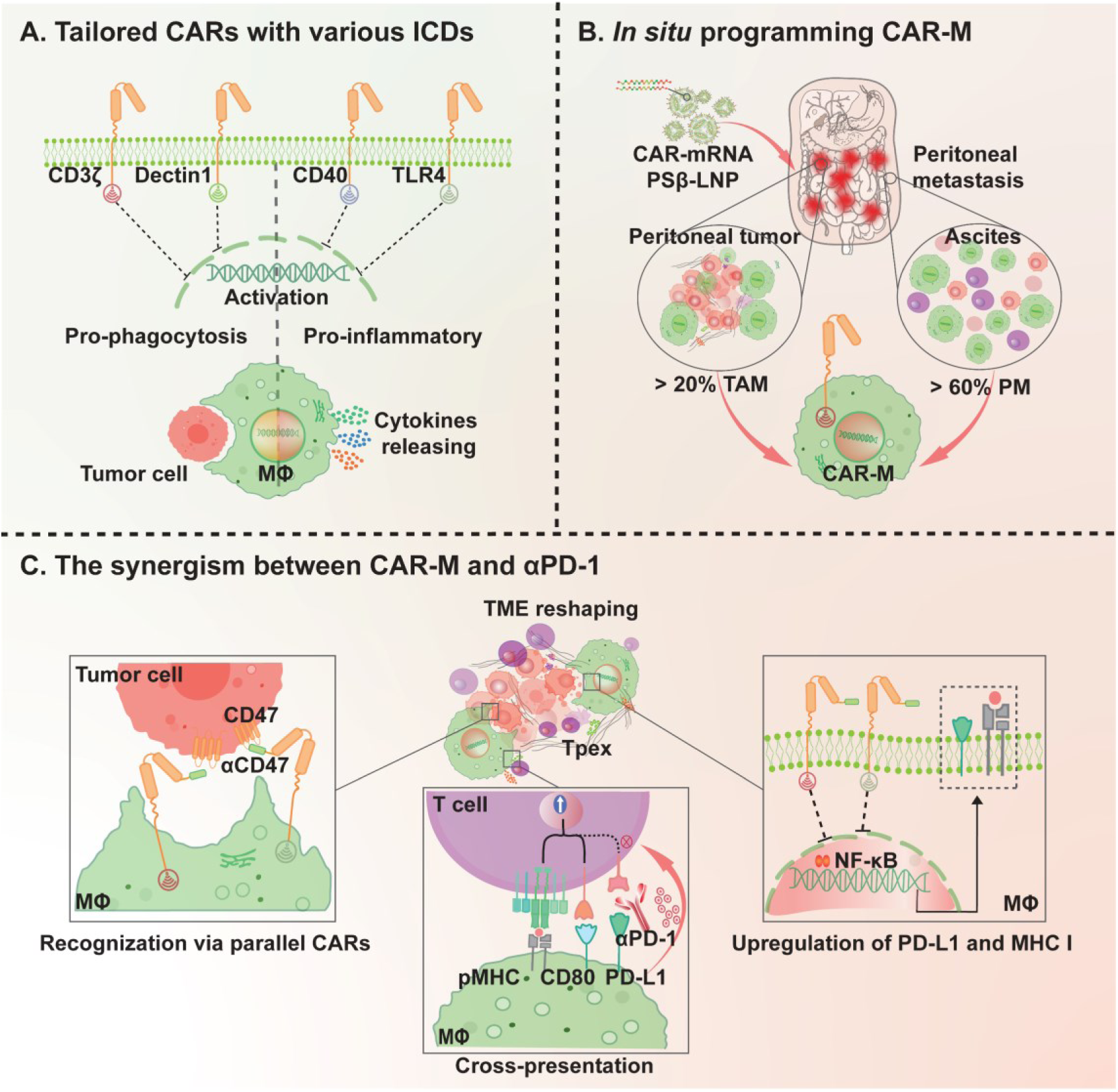
Schema of intraperitoneal programming of CAR-M via mRNA-LNP for reshaping TME and boosting cancer immunotherapy. mRNA, messenger RNA; CAR-M, chimeric antigen receptor macrophage; ICD, intracellular domain; LNP, lipid nanoparticle; TME, tumor microenvironment.

## Methods

### Cells and culture conditions

Cell lines THP-1, HS578T, MDA-MB-231, RAW264.7, 4T1 and CT26 used in this study were purchased from ATCC. SKOV3-luc, CT26-luc and ID8-luc were purchased from iCell. KPC-luc was a kind gift from the Penghui Zhang laboratory from HIM. eGFP and luciferase-containing GV260 lentiviral vectors (Ubi-eGFP-Firefly Luciferase-Ires Puromycin, Genechem) were transfected into HS578T and MDA-MB-231 for stable HS578T-luc and MDA-MB-231-luc cell lines as previously described. Human ERBB2 (HER2), eGFP, and luciferase-containing lentiviral vectors (Ubi-HER2-eGFP-Firefly Luciferase-IRES-Puromycin, Genechem) were transfected into CT26 and 4T1 for stable CT26-eGFP-luc-HER2 and 4T1-eGFP-luc-HER2 cell lines. All cell lines were cultured in complete RPMI-1640 with 10% origin FBS (Gibco), 1% penicillin/streptomycin (Gibco) and grown at 37 °C and 5% CO2 with saturating humidity. Bone marrow-derived macrophages (BMDMs) were generated as previously described with minor modification^57^. In brief, single-cell supernatants collected from murine femurs were cultured in complete RPMI-1640 with 20 ng/mL M-CSF (Sangon) for 7 days to harvest M0 macrophages and 20 ng/mL IL-4 (Sangon) for another 2 days to harvest M2a macrophages.

### mRNA construction and synthesis

The CARs in this study are composed of antigen-binding domain (scFv from Trastuzumab or mutated SIRPα), CD8α hinge, transmembrane domain, and intracellular domains from CD3ζ, CD40, CD46, CSF2R, Dectin1, and TLR4. Responding codon-optimized CAR plasmid was constructed with a T7 promoter, 5’-UTR and 3’-UTR, CAR sequence and poly (A)-tail. After linearization via restriction endonuclease BspQI, purified DNA was used as a template to transcribe CAR-mRNA via a T7 High Yield RNA Transcription Kit (Novoprotein). Notably, all uridines throughout the mRNA sequence were substituted with modified nucleobase N1-methyl-pseudouridines (m1ψ). IVT mRNA capped with a Cap 1 Capping System Kit (Novoprotein) was purified and diluted to 1 μg/μL for standby.

### Construction and characterization of mRNA-LNP

mRNAs diluted in citrate buffer (pH 4.0) were encapsulated in lipid nanoparticles (LNP) via self-assembly. Briefly, lipids, including ethanol containing ionizable cationic lipid (SM102), phosphatidylcholine (DSPC), cholesterol analogs (β-sitosterol), polyethylene glycol-lipid (DMG-PEG2000) and phosphatidylserine (DOPS) (molar ratio: 50:10:34:1:5), and mRNA in citrate buffer (pH 4.0) were mixed rapidly as volume ratio 1:2 and flow rate 20 mL/min via the microfluidic platform INano™ E (Micro&Nano). After pH balance and ultrafiltration, mixed mRNA-LNPs were stored at 4 °C for subsequent characterization and evaluation within 3 days. The physicochemical properties of RNA-loaded particles, including radius and polydispersity, were characterized by Zetasizer (Malvern). The morphology of particles was characterized by transmission electron microscope (TEM, JEM-2100plus from JEOL). Efficacy of the encapsulated particles was assessed via a Quant-iT™ RiboGreen™ RNA Assay Kit (Thermo Fisher). Theoretical calculation of pKa of particles was characterized via a TNS binding assay. Virtual lysosomal escape capability (membrane fusion and destabilization) was characterized via hemolysis assay at neutral and acidic pH, as described previously^58^.

### Transfection of macrophages

After being seeded in wells for several hours, macrophages (BMDM, THP-1, or RAW264.7) were cocultured with mRNA-LNP (typically 1 μg mRNA for 1.0 × 10^6^ cells) in complete medium at 37 °C. Transfection efficiency was analyzed using eGFP or CAR expression proportionally by flow cytometry (Beckman Coulter, CytoFLEX) and microscopy (Olympus ckx53). CARs were labeled with HER2 protein-conjugated Alexa Fluor 647.

Beads-based phagocytosis assay. HER2-PS beads were obtained from carboxylated polystyrene spheres (Bangs Laboratories) coupled with HER2 protein via crosslinkers 1-ethyl-3-(3- dimethylaminopropyl) carbodiimide hydrochloride (EDC) and N-hydroxy succinimide (NHS). 18 h after transfection, CAR-Ms were cocultured with PS-HER2 beads (ratio: 1:10) for 1 h at 37 °C. The uptake was stopped immediately by lowering the temperature. Beads-based phagocytosis was characterized via microscopy (TECAN). For quantitative analysis via flow cytometry (Beckman Coulter), HER2-PS beads were conjugated with fluorescein isothiocyanate (FITC), and macrophages were stained with the lipophilic membrane labeling probe 1,1’-dioctadecyl-3,3,3’,3’- tetramethylindocarbocyanine perchlorate (DIL) dye (Beyotime).

### *In vitro* cytotoxicity assay

After being plated in white flat non-transparent wells (Corning), BMDMs were transfected with the corresponding CAR-mRNA (1 μg mRNA into 1.0 million BMDMs). Following 18 hours of transfection, luciferase-loaded tumor cells were added in various E:T ratios, systematically adjusted from 10:1 to 1:1. After incubating for a total of 24 hours, bioluminescence luciferase substrate (D-Luciferin sodium salt, APExBIO) was added to achieve a final concentration of 1.5 mg/mL, and the luminescence signal was measured within 10 minutes using a microplate reader (TECAN Spark). For imaging with IVIS Spectrum, the white wells were replaced with black transparent wells (Corning). To image the engulfment of debris by macrophages, tumor cells were labeled with pHrodo^TM^ Red (Invitrogen™). For the phagocytosis process study, CAR-M and GFP-loaded tumor cells were monitored via Incucyte SX5 (Satorius), and images were captured every one and a half hours for three days.

### Cytokine analysis

After being plated in white flat non-transparent wells (Corning), BMDMs were transfected with the corresponding CAR-mRNA (1 μg mRNA into 1.0 million BMDMs). Following 18 hours of transfection, tumor cells were added proportionally at an E:T ratio of 5:1. After incubating for a total of 12 hours, the supernatant was collected from medium and subsequently diluted with 1× DPBS at different multiples. Secretion of cytokines (IL-6, IL-12, TNF-α) was tested using the corresponding ELISA assay kit (Sangon Biotech) and analyzed by a microplate reader (TECAN Spark).

### RNA sequencing of macrophages

After coculturing BMDM/CAR-Ms and targeted tumor cells or blank control for several hours, total RNA was extracted and isolated. The raw image data from sequencing results were processed using the Bcl2fastq software (v2.17.1.14) for base calling. Cutadapt (version 1.9.1) was used to preprocess the raw data, filtering out low-quality data and removing contamination and adapter sequences. Short reads were aligned using the Hisat2 (v2.0.1) software with default parameters. RNA-Seq data were analyzed for differential alternative splicing using rMATS (version 4.1.0). Based on the alignment results of each sample to the reference genome, the samtools (v0.1.19) software was used for mpileup processing to obtain potential SNV results for each sample. Subsequently, the annovar (v2016.05.11) software was used for annotation. Gene expression was calculated using HTSeq software (v0.6.1), which utilizes the FPKM (Fragments Per Kilobase per Million reads) method to quantify gene expression levels. We filtered the detected results according to the criteria of significant differential expression (fold change in gene expression of 2 or more and q-value ≤ 0.05).

### Mice and *in vivo* tumor models

All mouse studies complied with the humane and ethical treatment of experimental animals and were approved by the Hangzhou Institute of Medicine, Chinese Academy of Science Animal Center. All mice used in this study were obtained from the Zhejiang Experimental Animal Center. All mice were housed in a clean, pathogen-free, and humid environment at 25 °C with a typical humidity level of 50%, under a 12-hour dark/12-hour light cycle, with sufficient sterile water and food supply.

To model peritoneal metastasis CRC tumors, 2.0 × 10^5^ colorectal cancer tumor cells CT26- luc were intraperitoneally injected (i.p.) into Balb/c mice (4 to 6 weeks old). An ovarian tumor CDX model was built via inoculation of 3.0 × 10^6^ SKOV3 cells into immunodeficient nude mice. To model peritoneal metastasis in pancreatic tumors, 1.0 × 10^6^ PAN02-luc cells were intraperitoneally injected (i.p.) into C57BL/6 mice (4 to 6 weeks old).

To determine target and transfection efficacy, GFP-mRNA LNP or DIR-labeled LNP was injected several days after inoculation. The dosage used for testing was 10 μg of mRNA or an equivalent dose of LNP with sampling for analysis after 16 hours. Macrophage (F4/80^+^) uptake signal, transfection signal and distribution profile were characterized by GFP and DIR, respectively, via flow cytometry, IF, and IVIS.

For the antitumor efficacy study, CAR-mRNA PSβ-LNP was injected daily (5 μg/mouse) from the 3rd day after the inoculation of tumor cells, and tumor growth was monitored every three days by bioluminescence imaging (IVIS Lumina Series III, PerkinElmer) and analyzed using Living Image v4.4 (Caliper Life Sciences).

For the rechallenge study, 1.0 × 10^6^ tumor cells were injected (s.c.), and tumor growth was monitored every other day. The weight of mice was recorded every three days. At the experiment endpoint, mice were euthanized in accordance with laboratory animal guidelines.

For the tumor challenge experiment using splenic cell adoptive transfer from cured mice, 10 million splenic cells from cured mice were intravenously injected into naive mice one day before tumor implantation, followed by subcutaneous implantation of 1 million tumor cells. Tumor growth in mice was recorded every other day.

For the immune challenge experiment in an established TME, 1 million tumor cells were implanted (s.c.) in advance. On the seventh day after implantation, 10 million splenic cells from cured mice were adoptively transferred. Tumor growth in mice was then monitored every other day.

To evaluate the combined antitumor effects of PD-1 antibody and *in situ* programming of CAR- M therapy in the CT26-luc syngeneic model, intervention started on the 9th day after implantation of 2.0 × 10^5^ CT26-luc tumor cells (i.p.). The dosing regimen was as follows: PD-1 antibody was administered every three days at a dose of 100 μg per mouse per administration, totaling three administrations. CAR-mRNA PSβ-LNP was administered once daily with a total daily mRNA dose of 7.5 μg per mouse (comprising 5 μg of CD3ζCAR mRNA and 2.5 μg of TLR4 CAR mRNA), totaling seven administrations. The combination group received both CAR-mRNA treatment and PD-1 antibody treatment as noted above. Tumor growth was monitored every three days using IVIS. Intervention in the later-stage tumor model began on the 12th day to ensure an adequate supply of tumor samples for subsequent IF analysis, Luminex assay and scRNA sequencing.

To evaluate the combined effects in the PAN02-luc syngeneic model, the intervention started on the 8th day after implantation of 1.0 × 10^6^ PAN02-luc tumor cells (i.p.). Intervention and monitoring measures were identical to those described above.

To evaluate the combined antitumor effects of siRNA-mediated macrophage PD-L1 knockdown and *in situ* programming of CAR M therapy, PD-L1 siRNA was administered starting on the 8th day after tumor implantation at a dose of 5 μg per mouse every three days, totaling three administrations, and CAR-mRNA PSβ-LNP was administered starting on the ninth day with a total mRNA dose of 7.5 μg per mouse per day (comprising 5 μg of CD3ζ CAR mRNA and 2.5 μg of TLR4 CAR mRNA), totaling seven administrations. The combination group received both of CAR-mRNA treatment and PD-L1 siRNA treatment as noted above. Tumor growth was monitored every three days using IVIS.

### TME study

Peritoneal ascites were collected for the tumor microenvironment study, and tumors were resected for immunohistochemical analysis and then digested for flow cytometry after red blood cell lysis. Immune cell panels were stained with anti-mouse flow cytometry antibody probes. Macrophage polarization was checked via M1 or M2 markers after intervention. Primary antibodies used in the study are listed as follows: F4/80 (123117, BioLegend), CD80 (104715, BioLegend), CD80 (50446-R014-A-25, Sino Biological), CD86 (105005, BioLegend), CD86 (50068-RP02-50, Sino Biological), CD206 (FAB2535P, R&D), CD206 (#24595, CST), CD3 (100228, BD), CD4 (550954, BD), CD8 (K0227-A64, MBL), IFNr (505807, BioLegend), and goat anti-rabbit IgG H&L (Alexa Fluor® 488) (ab150077, abcam). All collected data were analyzed using FlowJo 10.8.1 software.

In addition, the secretion of multiple cytokines was tested via a Luminex assay. The hierarchical clustering plot was generated using the R software (v.4.2.2) package ape (v.5.6.2)^59^ through Hiplot Pro (https://hiplot.com.cn/), a comprehensive web service for biomedical data analysis and visualization.

### T cell activation and IFNγ ELISPOT

T cells collected from ascites and tumors were stained with flow cytometry antibody probes. Activated CD8^+^ T cells were determined as CD3^+^CD8^+^IFNr^+^, and TH1 cells were determined as CD3^+^CD4^+^IFNγ^+^. For the rechallenge study, an ELISPOT assay was performed to evaluate T-cell activation. In summary, 1.5 × 10^5^ splenocytes collected from cured mice or naïve mice after RBC lysis were seeded in wells of a 96-well IFNγ ELISPOT plate with 1.5 × 10^4^ tumor cells to stimulate tumor-specific T cells. After incubation for 18 hours, the ELISpot assay was conducted utilizing the IFN-γ ELISpot kit (ThermoFisher) in accordance with the manufacturer’s instructions. Spots were captured under an automated ELISpot and FluoroSpot reader (Mabtech Astor2).

### Tumor and spleen digestion

The excised tumors and spleens were cut into small pieces of approximately 3 mm^3^. Then, these small pieces were incubated in a Collagenase/Hyaluronidase (10X collagenase/hyaluronidase in DMEM, STEMCELL) solution for one hour before passing through a 300-mesh cell strainer. The targeted single-cell population was obtained after RBC lysis and centrifugation.

### Single-cell RNA-seq

Single-cell counting and quality control were performed using the TC20 Automated Cell Counter (Bio-Rad, USA). Then, after GEM (Gel Bead-in-emulsion) generation, 10x barcoded, full-length cDNA amplification was obtained from polyadenylated mRNA. After library construction, all repositories with different indices were multiplexed and loaded on an Illumina NovaSeq instrument according to the manufacturer’s instructions (Illumina, San Diego, CA, USA). Raw sequencing data quality control and mapping were performed with a 10X genomics single-cell gene expression analysis pipeline. Subsequently, cell quality control, clustering, and marker gene analysis were performed using Seurat (v4.3.0). We excluded genes expressed in extremely few cells to maintain the quality of genes and cells. Then, SingleR (v2.0.0) was utilized for cell type annotation. GOSeq (v1.34.1) was employed for Gene Ontology (GO) enrichment analysis, in-house scripts were used for KEGG (Kyoto Encyclopedia of Genes and Genomes) enrichment analysis, reactomPA (v1.42.0) was used for reactome pathway enrichment, and GSVA (v1.46.0) was applied for gene set variation analysis (GSVA). Additionally, Monocle was utilized to infer the pseudotime trajectory of cells.

All software used for single-cell RNA analysis in the study was listed as follows:

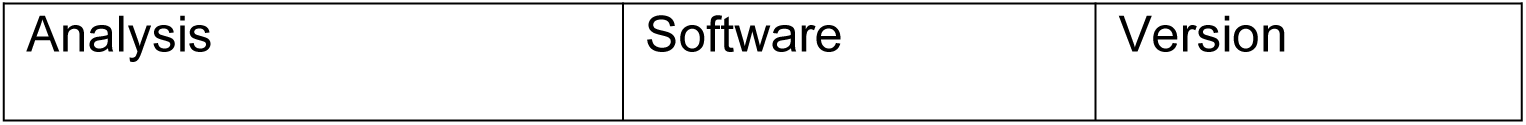
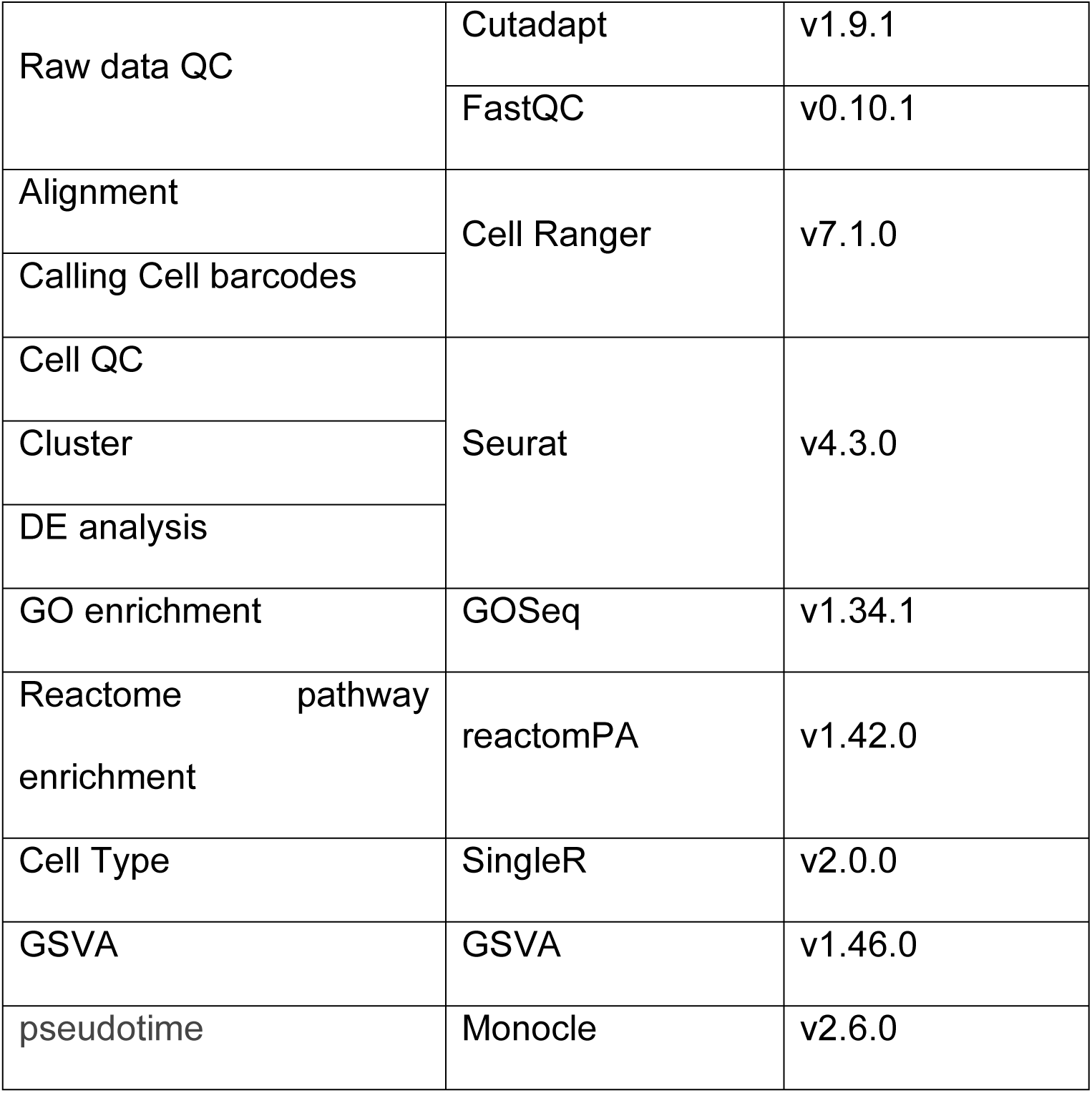

### Biocompatibility

To evaluate biosafety, 7.5 ug CAR-mRNA PSβ-LNP or an equivalent dose of empty LNP was injected intraperitoneally into healthy Balb/c mice daily. After 7 consecutive days, the mice were euthanized. Primary organs, including heart, liver, lung, kidney, spleen, and intestine, were collected for H&E staining and pathological analysis, and corresponding blood was collected for blood biochemistry and routine blood tests. Blood counts were performed via an automatic hematology analyzer (BC-2800vet, Mindary).

## Statistics

All data analyses were conducted using GraphPad Prism (v 8.0) with results presented as mean ± SD. P-values were determined using one/two-way analysis of variance (ANOVA) for tumor growth, followed by multiple comparison tests as indicated in the figure captions. Spearman’s rho correlation test was used to assess correlations, a log-rank test was used for survival analysis, and unpaired two-tailed t-tests were used for other analyses. A P-value less than 0.05 was considered statistically significant. Experiments were repeated multiple times as independent trials.

## Supporting information

SI-Intraperitoneal programming of tailored CAR macrophages via mRNA-LNP to boost cancer immunotherapy

## Acknowledgments

This work is supported by the National Key Research and Development Program of China (2022YFC3401402 and 2023YFC3405100), the National Natural Science Foundation of China (NSFC T2188102, 22104133 22274141 and 32201143) and the Zhejiang Provincial Natural Science Foundation of China (LDQ23B050001 and LDQ24B020002). Authors acknowledge support from the Shared Instrumentation Core Facility, Hangzhou Institute of Medicine (HIM), Chinese Academy of Sciences.

## Author contributions

K.G., X.L., S.X. and W.T. contributed to conceptualizing the research project, experimental design, data curation and data analysis. S.X. and W.T. secured the funding for the study. T.L., L.H., Y.Z., W.Y., M.Z., Y.C., H.W., M.W. and Y. Z. contributed to partial experimental implementation. B.L., S.L., Y.Z. and J.S. contributed to data curation and data analysis. Y.Z., C.Z., W.W., Z.D. and P.Z. contributed to the experimental design. The manuscript was written by K.G. with revisions made by X.L., S.X. and W.T.

## Additional information

The online version contains supplementary material, including Supplementary Fig. 1-19, Table 1- 3, and Movie 1 and 2.

