## Supplementary material for "Intraperitoneal programming of tailored CAR macrophages via mRNA-LNP to boost cancer immunotherapy": SI-Intraperitoneal programming of tailored CAR macrophages via mRNA-LNP to boost cancer immunotherapy

**The supplementary information includes:**

**Supplementary Fig. 1-19**

**Supplementary Table 1-3**

**Other supplementary information:**

**Supplementary** **Movie 1 and 2**

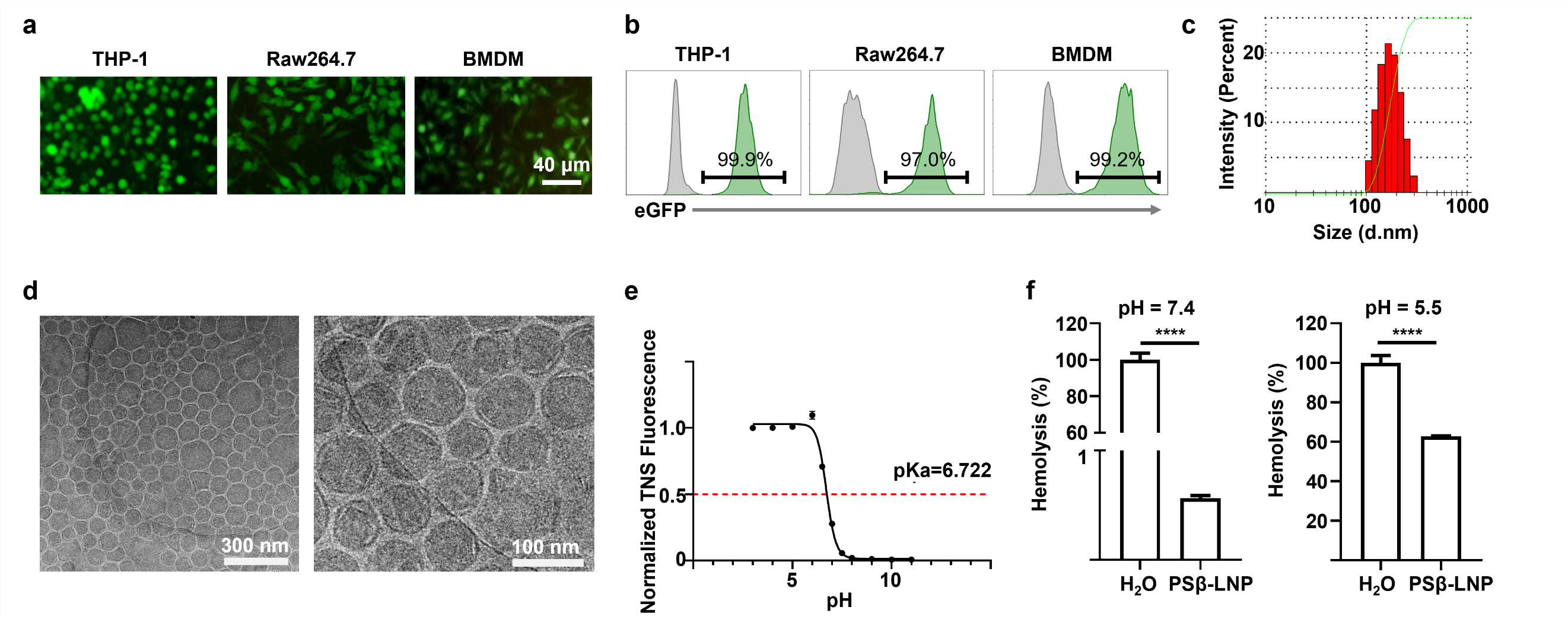

**Supplementary Fig. 1 |** Transfection profiles and physicochemical properties of the PSβ-LNP. Microscopic fluorescence imaging **(a)** and flow cytometric analysis **(b)** of macrophages transfected with the eGFP mRNA-encapsulated PSβ-LNP**. (c)** The size distribution of the PSβ-LNP was measured by the dynamic light scattering (DLS). **(d)** TEM imaging of PSβ-LNP. **(e)** The pKa of PSβ-LNP was measured via the 2-(p-toluidino) naphthalene-6-sulfonic acid (TNS) fluorescence method. **(f)** Hemolysis analysis of the PSβ-LNP in pH 7.4 and pH 5.5 conditions. Data are shown as mean ± SD (n = 3 independent duplicate samples). Statistical significance was calculated using unpaired *t*-test. For all panels, ns = no significance, **P* < 0.05, ***P* < 0.01, ****P* < 0.001, *****P* < 0.0001.

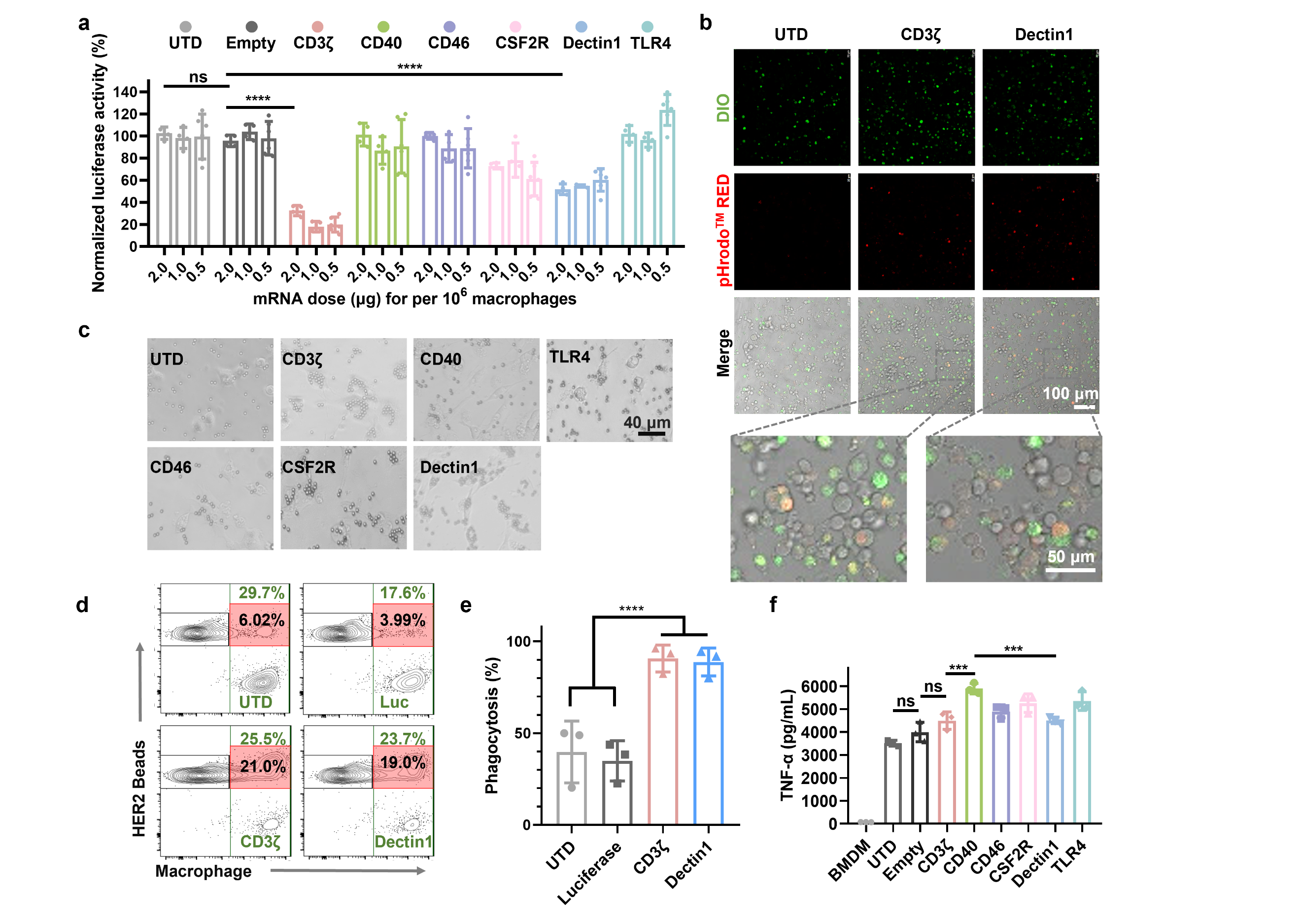

**Supplementary Fig. 2 |** **(a)** The normalized analysis on phagocytosis of luciferase-reported tumor cells by anti-HER2 CAR-Ms with pending ICDs in a series of mRNA doses (2.0 μg, 1.0 μg and 0.5 μg, respectively for 10^6^ macrophages). Data are shown in means ± SD (n ≥ 4). Statistical significance was calculated using two-way ANOVA. **(b)** Confocal images of CAR-M (green) phagocyting tumor cells stained with pHrodo^TM^ red. **(c)** Images of CAR-M or ctrl phagocyting HER2-conjugated beads. Flow cytometry analysis **(d)** and statistic **(e)** of CAR-M or ctrl (DID stained) phagocyting HER2 protein/FITC-conjugated beads. Data are shown in means ± SD (n = 3). Statistical significance was calculated using one-way ANOVA. **(f)** TNF-α analysis in the medium supernatant of controls or CAR-Ms after 24 h coincubation with SKOV3 tumor cells. Data are shown in mean ± SD (n = 3). Statistical significance was calculated using one-way ANOVA. For all panels, ns = no significance, **P* < 0.05, ***P* < 0.01, ****P* < 0.001, *****P* < 0.0001.

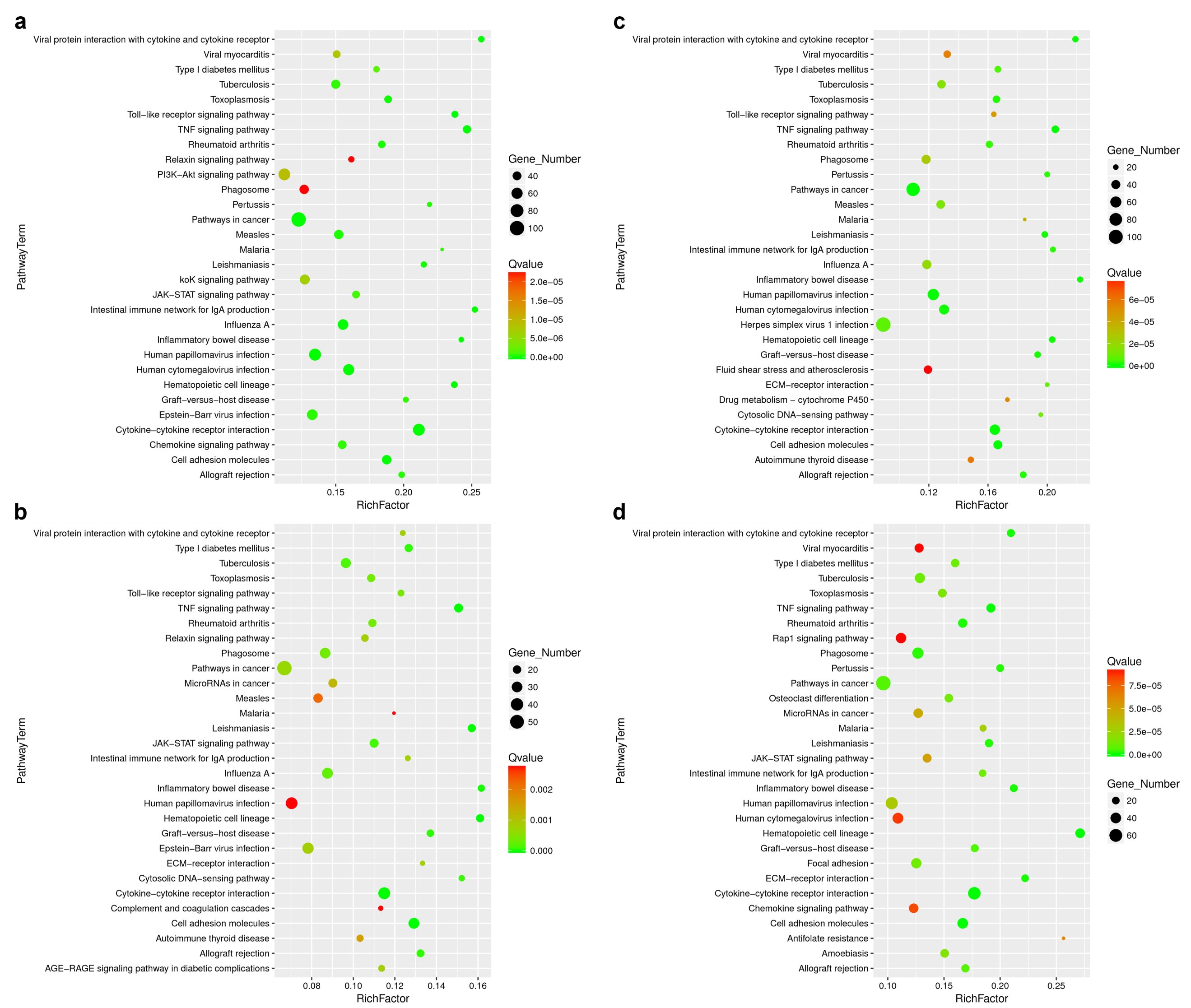

**Supplementary Fig. 3 |** Kyoto Encyclopedia of Genes and Genomes (KEGG) analysis (**(a)** CD3ζ vs Empty; **(b)** CD40 vs Empty; **(c)** Dectin1 vs Empty; **(d)** TLR4 vs Empty) of macrophages after various CARs signal transduction.

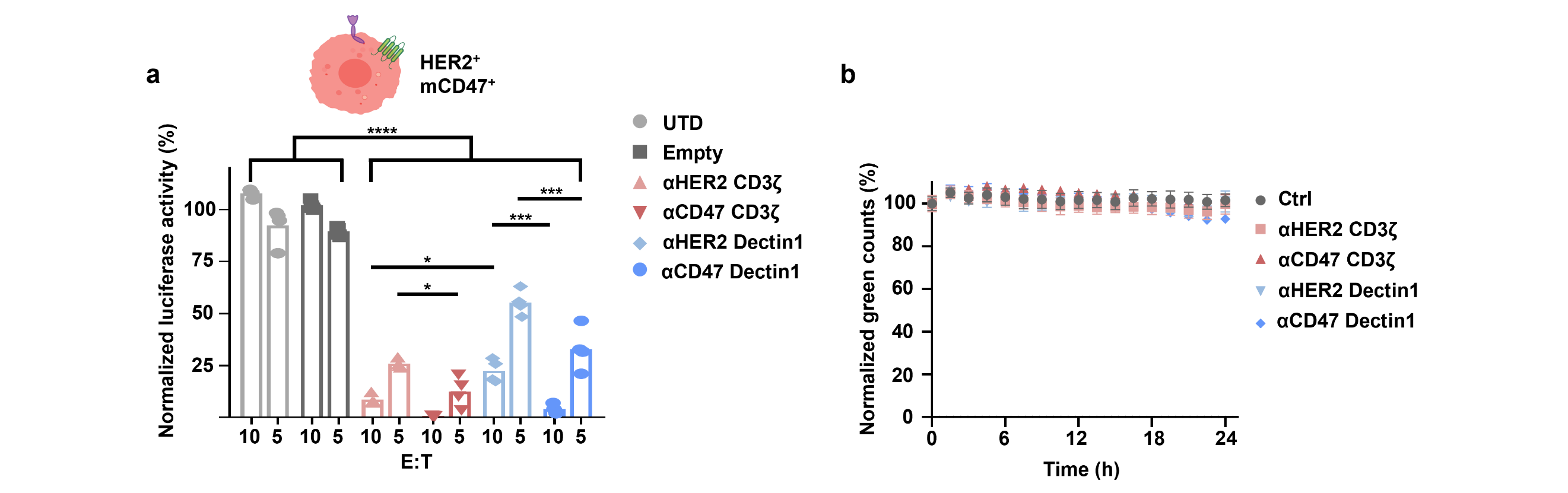

**Supplementary Fig. 4 |** **(a)** The normalized analysis on phagocytosis of luciferase-reported tumor cells (4T1-huERBB2^+^-mCD47^+^) by CAR-Ms. Data are shown as mean ± SD (n = 3 independent duplicate samples). Statistical significance was calculated using two-way ANOVA. **(b)** Incucyte-based phagocytosis assay of MDB231 (huERBB2^−^-huCD47^−^-eGFP^+^) by CAR-Ms or control within 24 hours. Data are shown as mean ± SD (n = 3 independent duplicate samples). For all panels, ns = no significance, **P* < 0.05, ***P* < 0.01, ****P* < 0.001, *****P* < 0.0001.

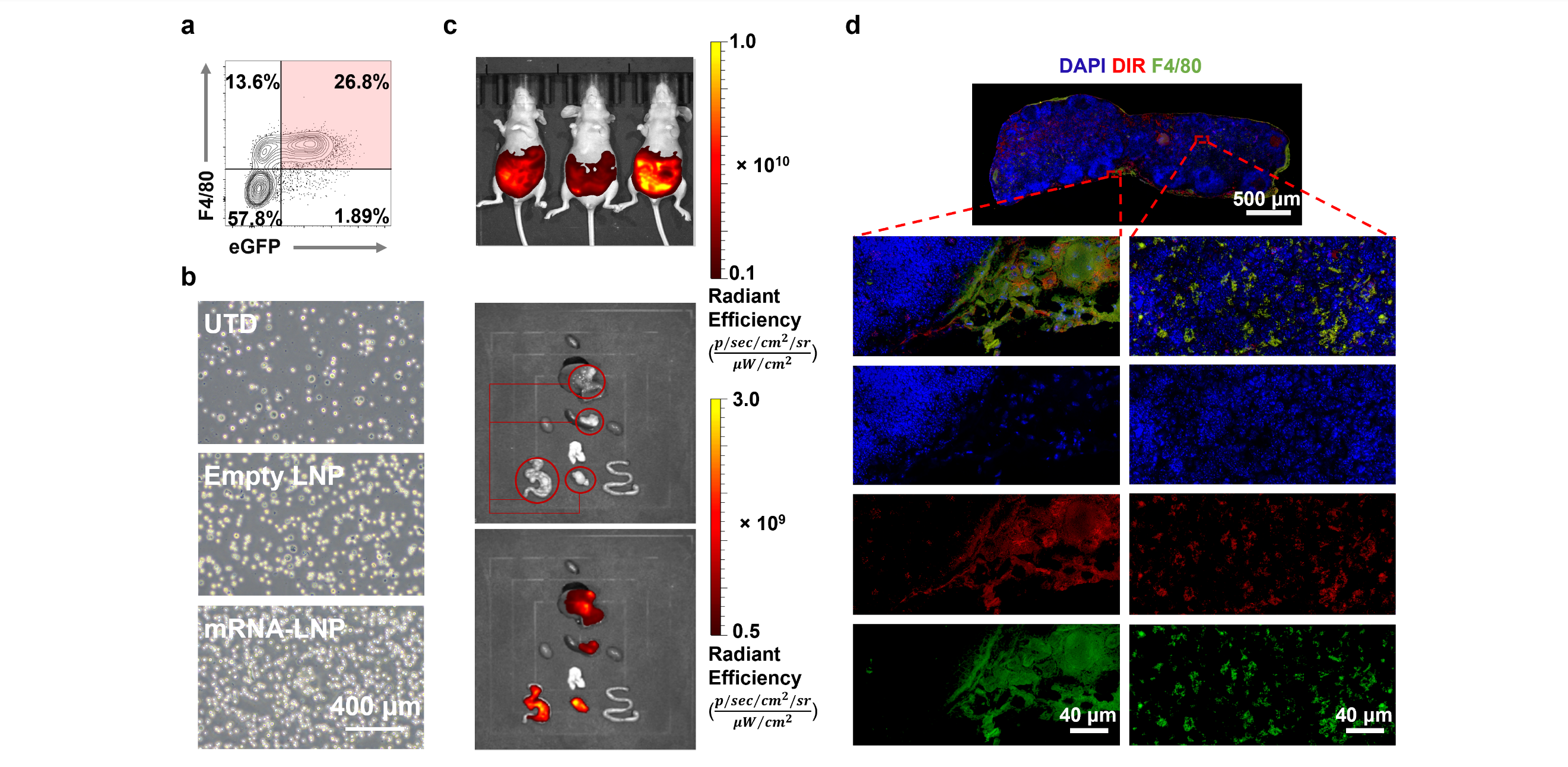

**Supplementary Fig. 5 |** **(a)** Flow cytometry analysis of eGFP expression proportion of ascites-derived cells after infusion of eGFP mRNA-PSβ-LNP in the PAN02-luc syngeneic mouse model. **(b)** Microscopic imaging of harvested peritoneal fluid cells, standardized to an identical dilution factor, following respective treatments on the 7th day after intraperitoneal implantation of 1 million CT26-luc cells. Scale bar, 400 μm. **(c)** DIR signal distribution after intraperitoneal administration of DIR labled PSβ-LNP in the SKOV3 xenograft mouse model. **(d)** Immunofluorescence slice (F4/80, green; DIR, red; nuclei, blue) of tumor tissue after DIR-PSβ-LNP infusion in the SKOV3 xenograft mouse model. Scale bar, 40 μm.

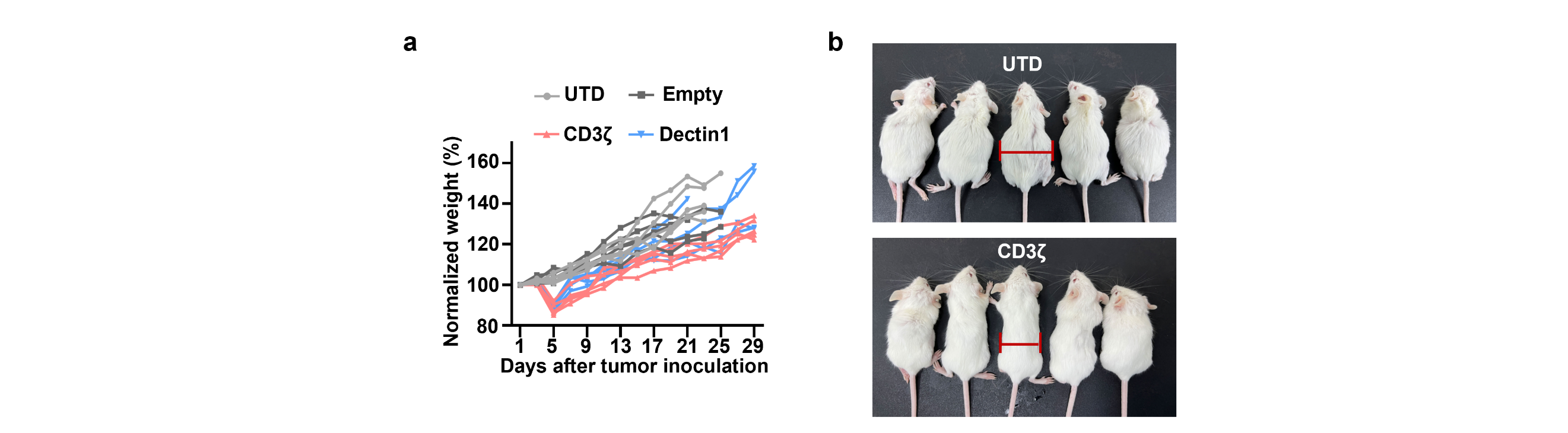

**Supplementary Fig. 6 |** **(a)** Normalized body weight of mice treated with the CD3ζ or Dectin1 CAR-mRNA PSβ-LNP or empty LNP in the CT26-luc syngeneic mouse model. **(b)** Representative images of the CT26-luc syngeneic mouse model.

­­
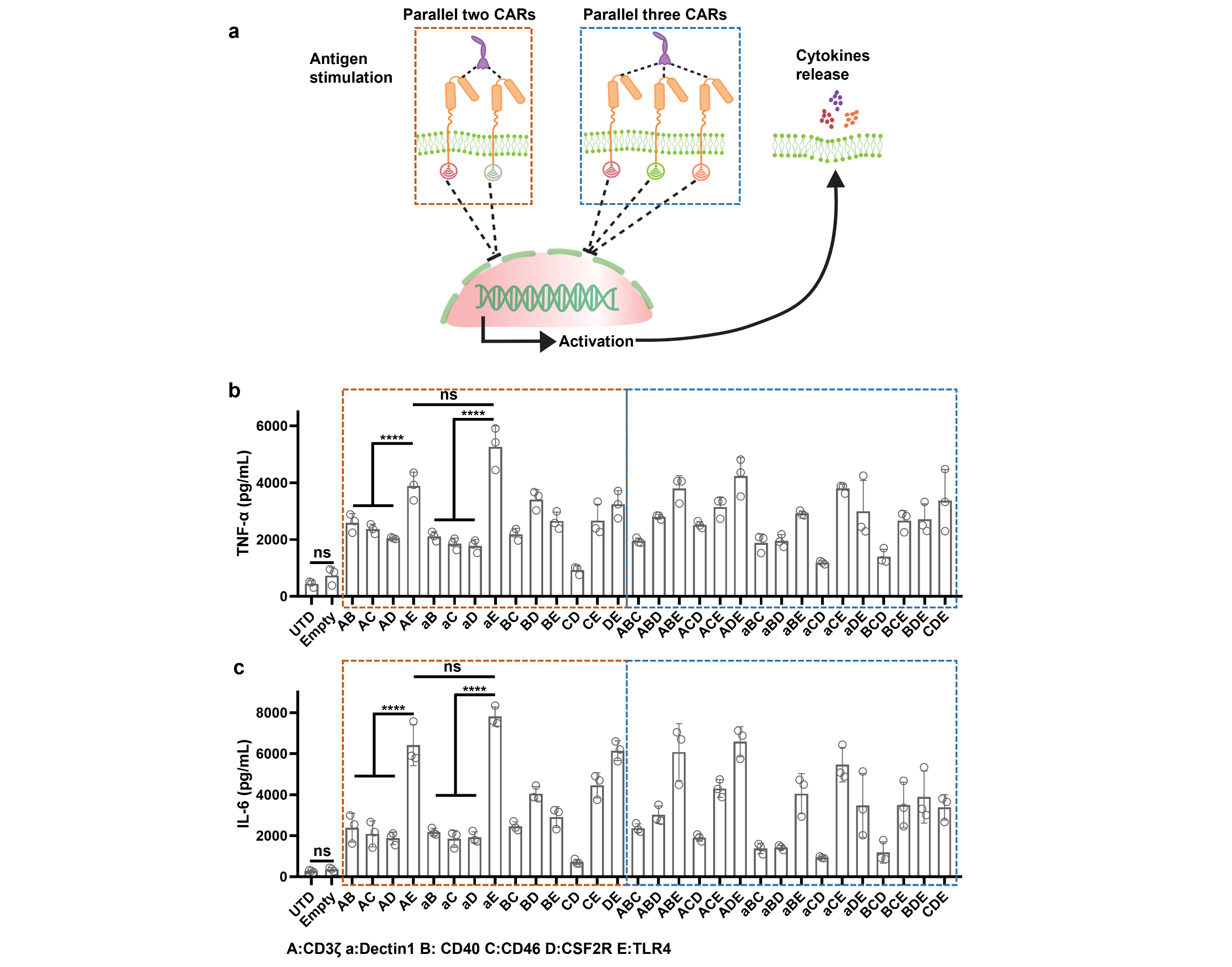

**Supplementary Fig. 7 |** **(a)** Schematic illustration of cytokine release induced via mono CAR or parallel CARs signal transduction. IL-6 **(b)** and TNF-α **(c)** were measured via ELISA in the groups. Data are shown as mean ± SD (n = 3 independent duplicate samples). Statistical significance was calculated using one-way ANOVA. For all panels, ns = no significance, **P* < 0.05, ***P* < 0.01, ****P* < 0.001, *****P* < 0.0001.

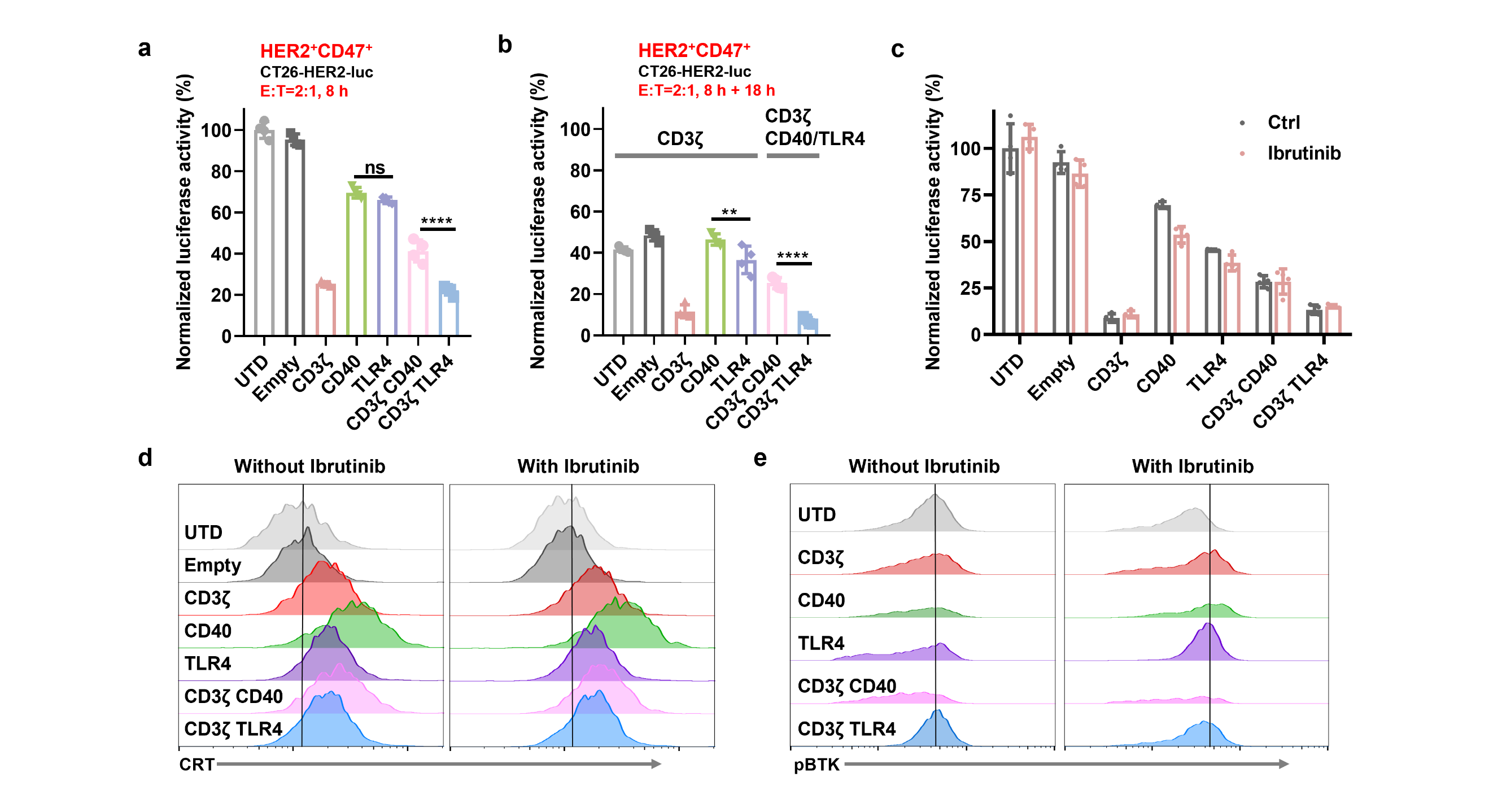

**Supplementary Fig. 8 |** **(a)** The normalized analysis on phagocytosis of luciferase-reported tumor cells (CT26-huERBB2^+^-mCD47^+^) by CAR-Ms for 8 h. E:T = 2:1. **(b)** Normalized phagocytosis after the second CD3ζ CAR mRNA transfection towards macrophages in **(a)** for another 18 h. For **(a)** and **(b)**, statistical significance was calculated using a one-way ANOVA. **(c)** Normalized phagocytosis of CT26-luc by CAR-Ms for 18 h with or without the Ibrutinib treatment. Data are shown as mean ± SD (n = 3 independent duplicate samples). Statistical significance was calculated using two-way ANOVA. Flow cytometry analysis of Calreticulin (CRT) **(d)** and phospho-Bruton's tyrosine kinase (pBTK) **(e)** expression with or without Ibrutinib treatment. For all panels, ns = no significance, **P* < 0.05, ***P* < 0.01, ****P* < 0.001, *****P* < 0.0001.

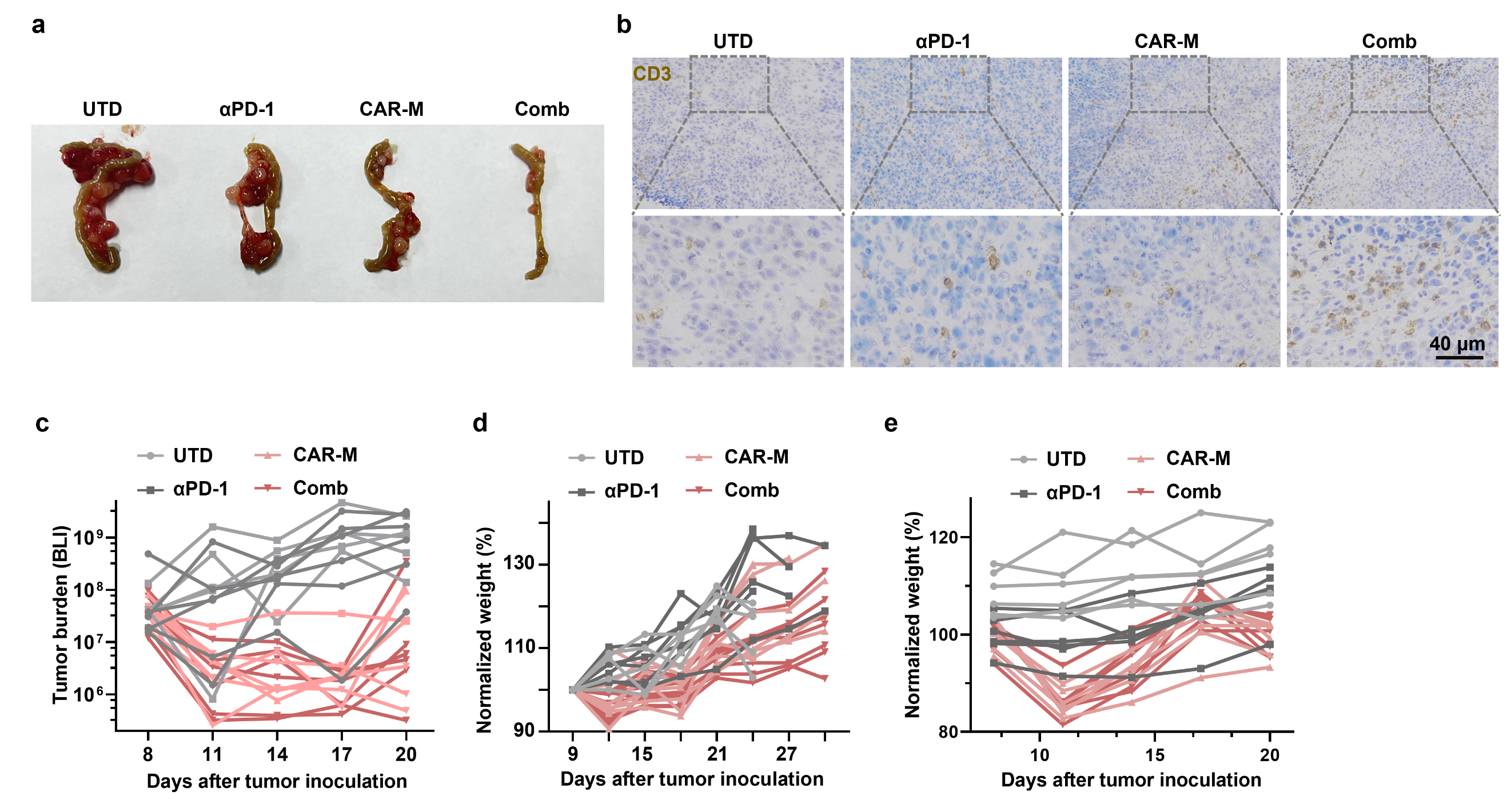

**Supplementary Fig. 9 |** **(a)** Representative images of tumors from the CT26-luc syngeneic mouse model. **(b)** Immunohistochemistry images of tumor tissues in the CT26-luc syngeneic mouse model. Scale bar, 40 μm. **(c)** Quantified BLI signal intensity of the PAN02-luc syngeneic mouse model with each treatment. **(d)** Normalized body weight of mice in cohorts of the CT26-luc syngeneic mouse model with each treatment. **(e)** Normalized weight of mice in cohorts of the PAN02-luc syngeneic mouse model with each treatment.

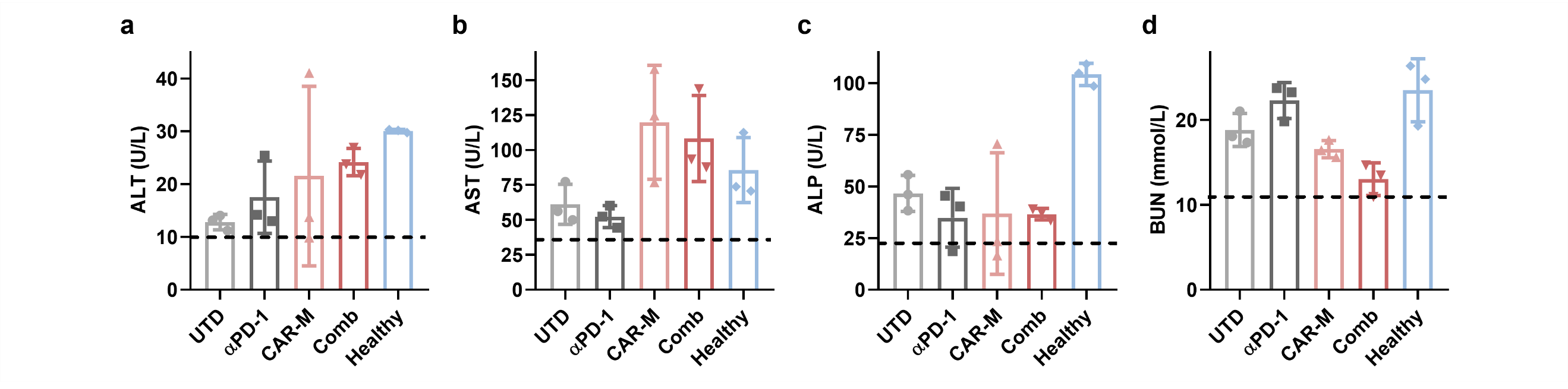

**Supplementary Fig. 10 |** Serum chemistry analysis (**(a)** Alanine Aminotransferase, ALT; **(b)** Aspartate Aminotransferase, AST; **(c)** Alkaline Phosphatase, ALP; **(d)** Blood Urea Nitrogen, BUN) of the CT26-luc syngeneic mouse model after the respective intervention. The black dashed line represents the lower limit of the safety threshold. Data are shown as mean ± SD (n = 3 independent duplicate samples).

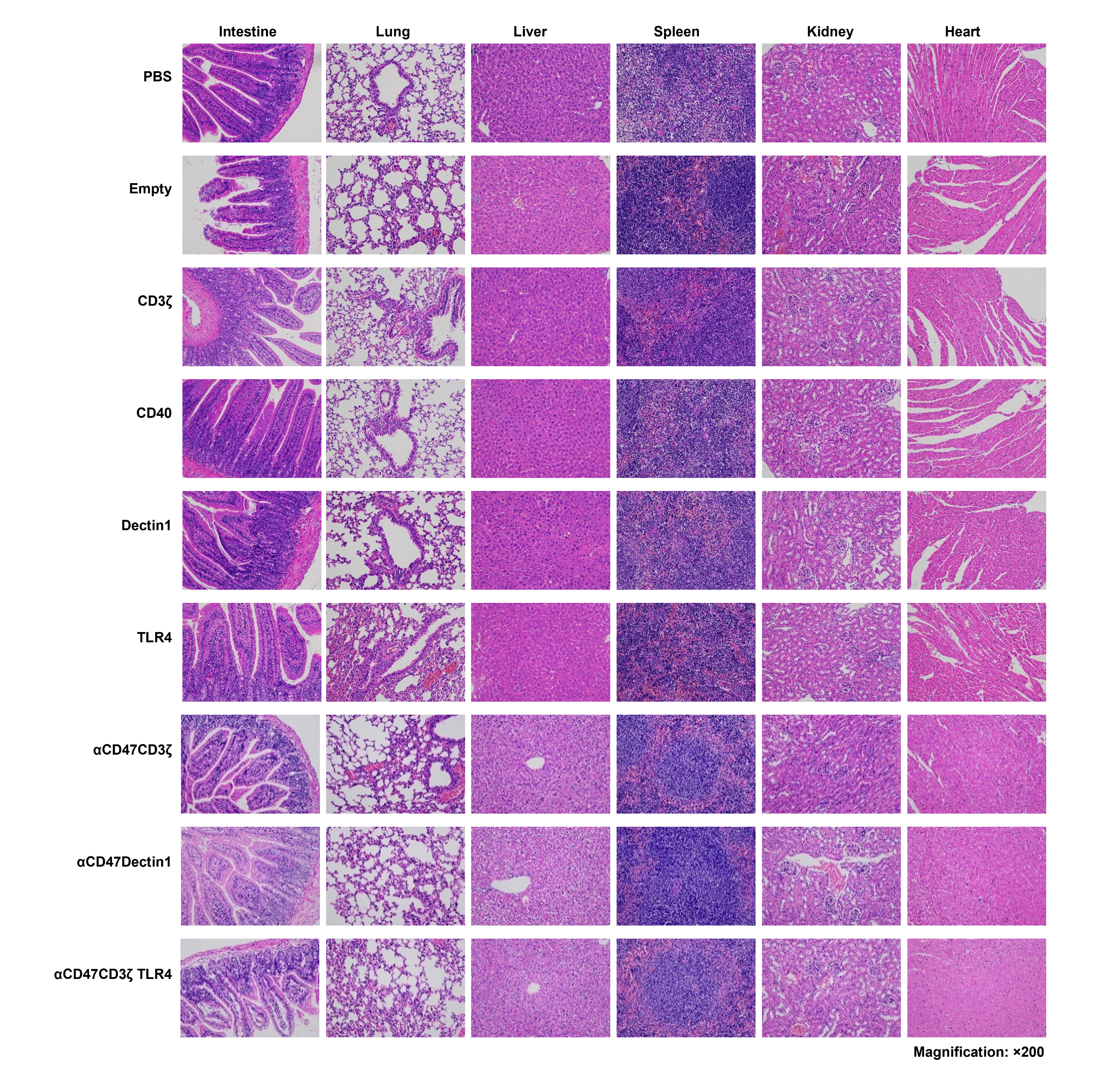

**Supplementary Fig. 11 |** H&E staining images of major organs in healthy female Balb/c mice under each intervention.

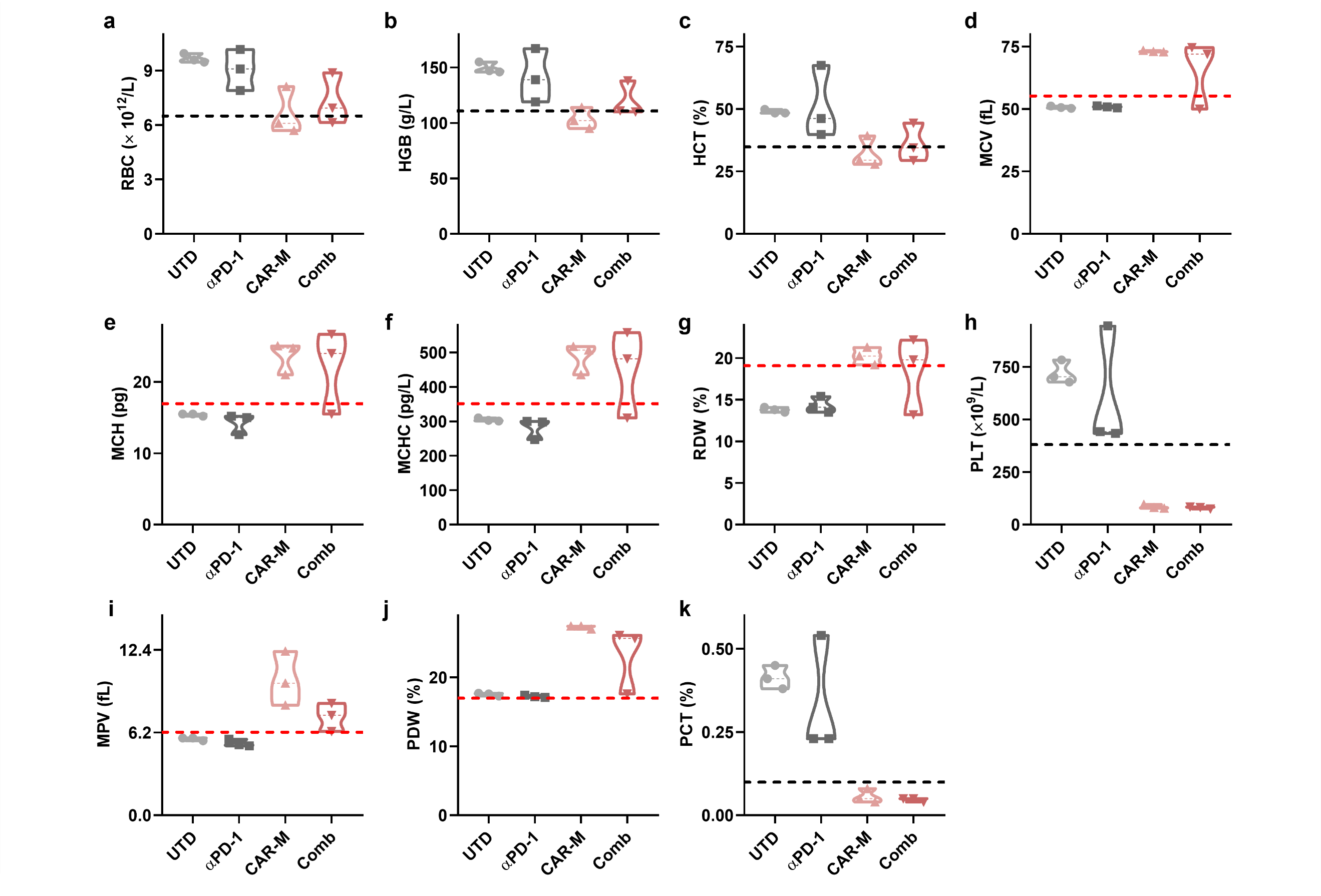

**Supplementary Fig. 12 |** Blood cells analysis of CT26-luc syngeneic mouse model after the respective intervention. (**(a)** Red Blood Cell, RBC; **(b)** Hemoglobin, HGB; **(c)** Hematocrit, HCT; **(d)** Mean Corpuscular Volume, MCV; **(e)** Mean Corpuscular Hemoglobin, MCH; **(f)** Mean Corpuscular Hemoglobin Concentration, MCHC; **(g)** Red Cell Distribution Width, RDW; **(h)** Platelet Count, PLT; **(i)** Mean Platelet Volume, MPV; **(j)** Platelet Distribution Width, PDW; **(k)** Plateletcrit, PCT)**.** The red dashed line and the black dashed line, respectively, represent the upper limit and the lower limit of the normal range. All data points are shown as violin plots (n = 3).

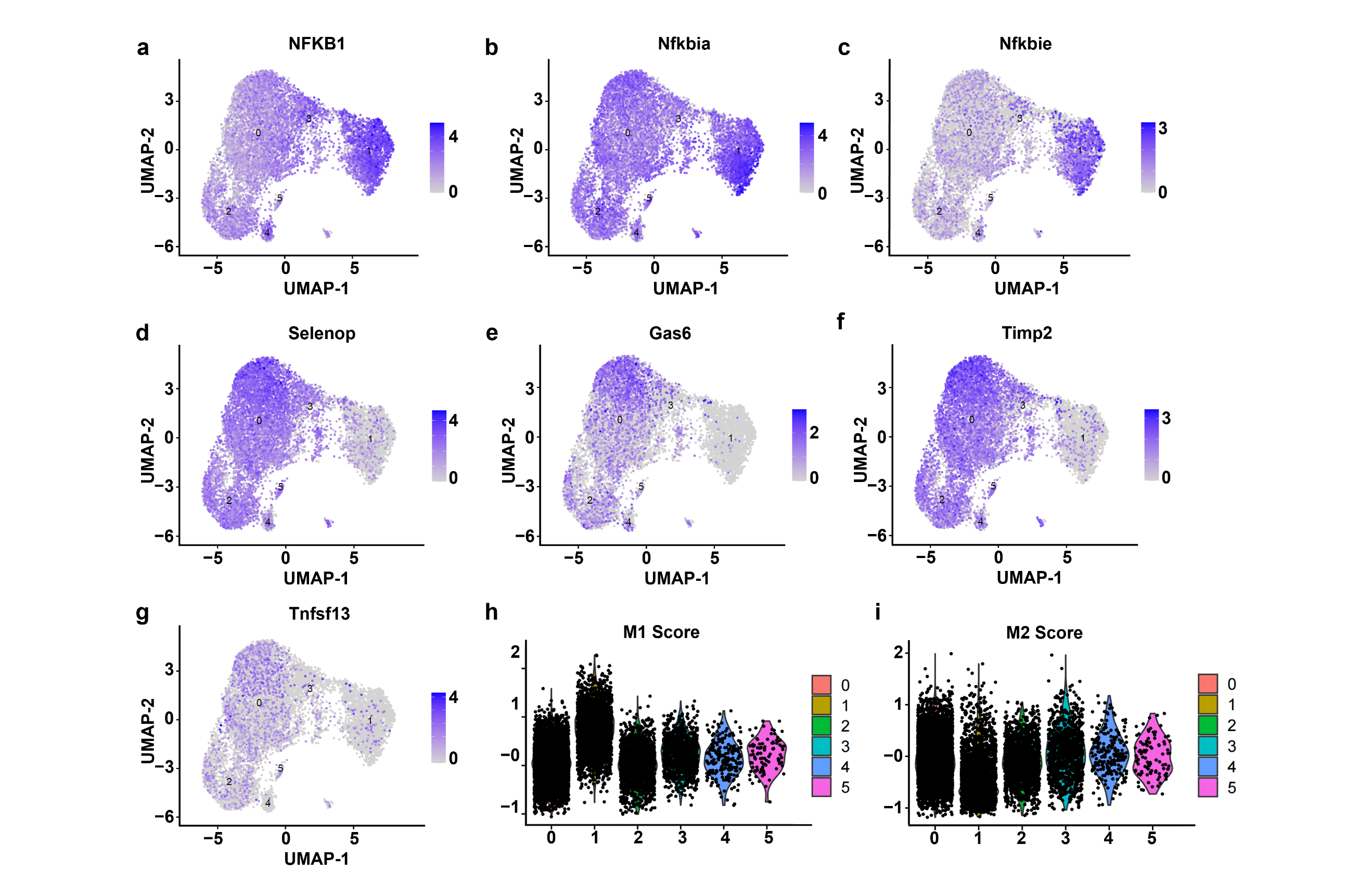

**Supplementary Fig. 13 |** UMAP plot of relative expression on indicated clustered macrophage genes. **(a)** NFKB1; **(b)** Nfkbia; **(c)** Nfkbie; **(d)** Selenop; **(e)** Gas6; **(f)** Timp2; **(g)** Tnfsf13. M1 score **(h)** or M2 **(i)** score of featured genes of clustered tumor-infiltrating macrophages.

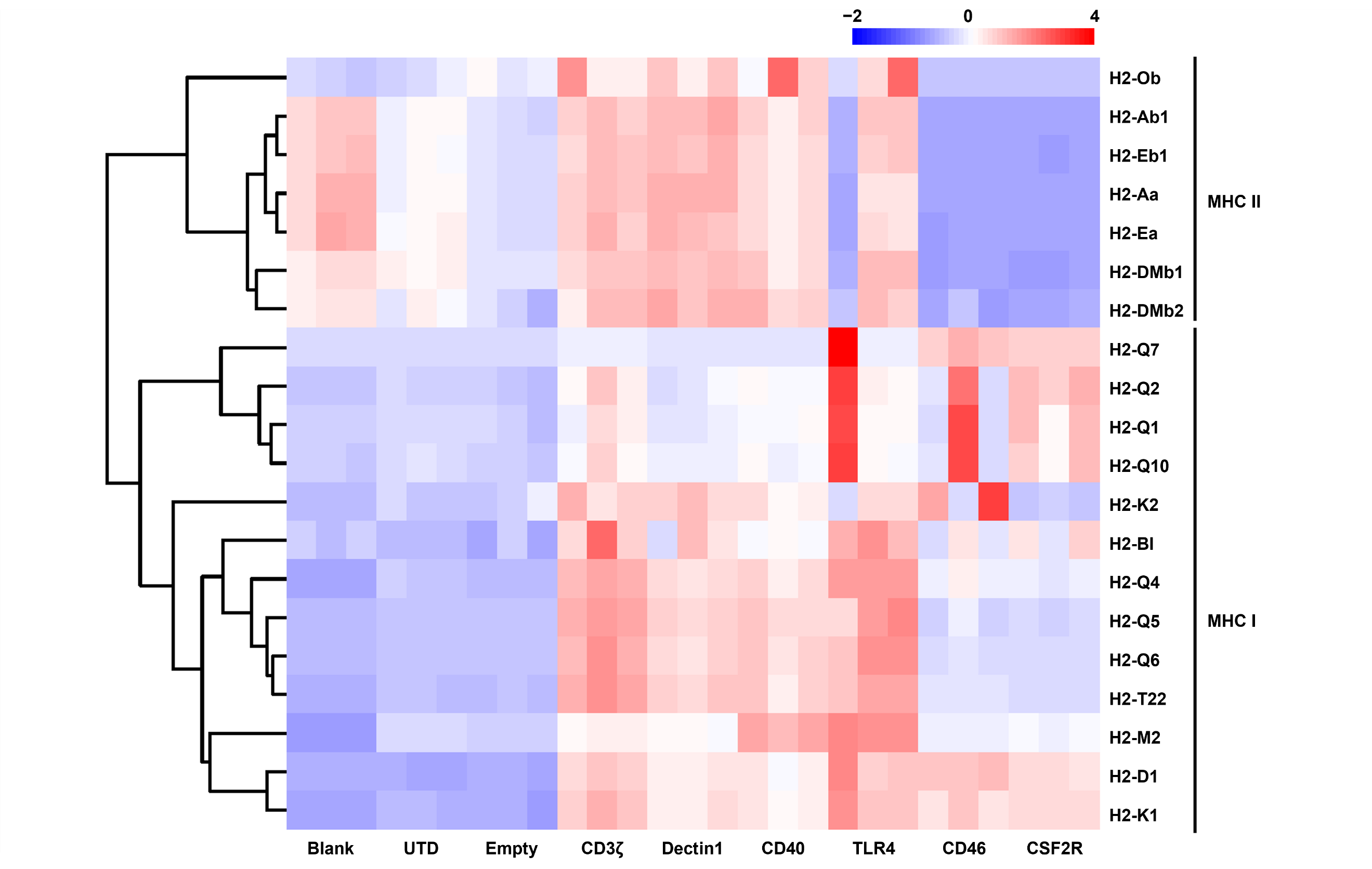

**Supplementary Fig. 14 |** Heatmap of MHC I and MHC II encoded gene expression in BMDMs after various CAR signal transduction.

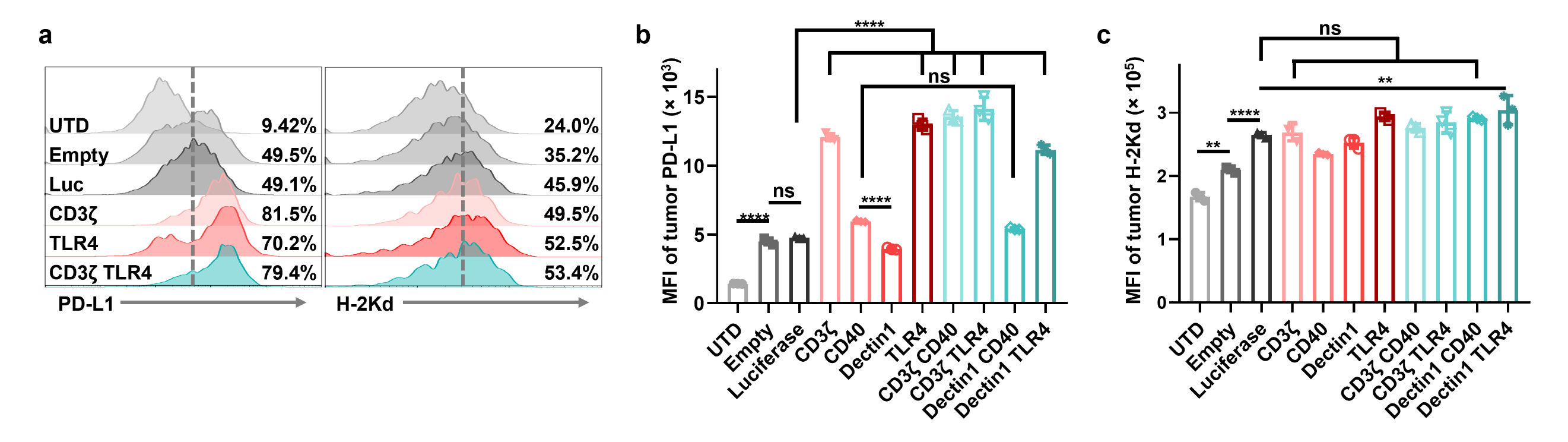

**Supplementary Fig. 15 | (a)** Flow cytometry analysis of PD-L1 and H-2Kd expression of tumor cells after interaction with CAR-M or control. Mean fluorescence intensity (MFI) was calculated from flow cytometry profiles on tumor PD-L1 **(b)** and H-2Kd **(c)** expression. Data are shown as mean ± SD (n = 3 independent duplicate samples). Statistical significance was calculated using one-way ANOVA. For all panels, ns = no significance, **P* < 0.05, ***P* < 0.01, ****P* < 0.001, *****P* < 0.0001.

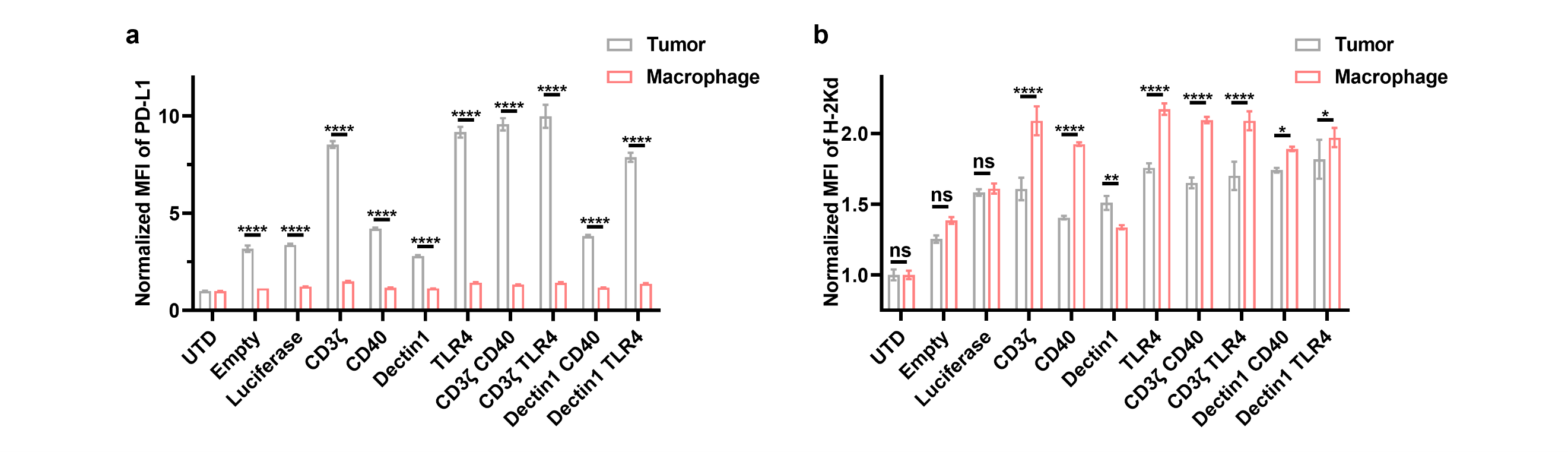

**Supplementary Fig. 16 |** Normalized PD-L1 **(a)** and H-2Kd **(b)** expression analysis on macrophages and tumor cells, respectively. Data are shown as mean ± SD (n = 3 independent duplicate samples). Statistical significance was calculated using two-way ANOVA. For all panels, ns = no significance, **P* < 0.05, ***P* < 0.01, ****P* < 0.001, *****P* < 0.0001.

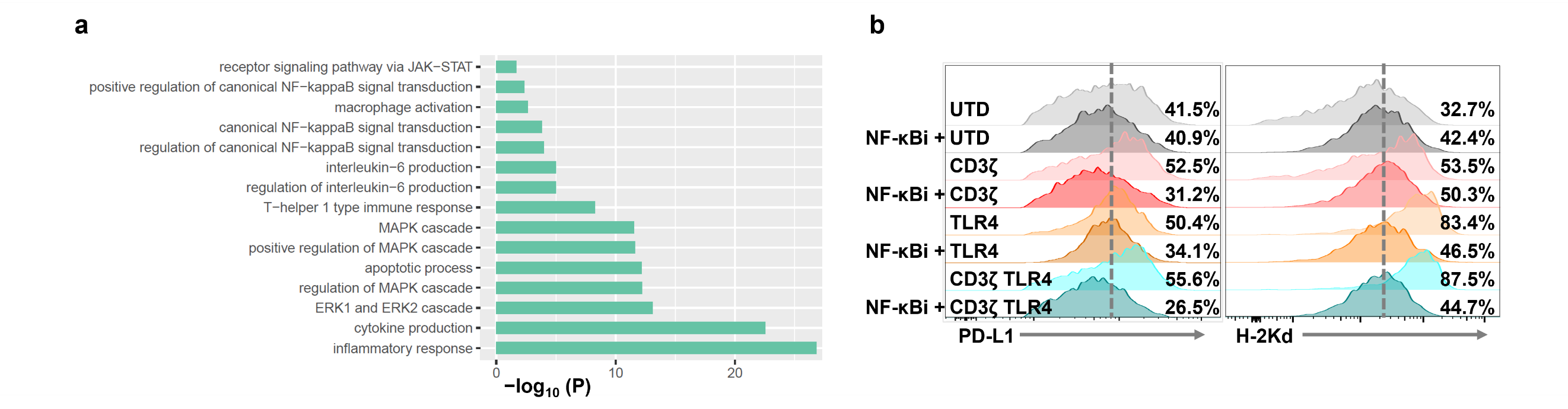

**Supplementary Fig. 17 | (a)** Gene Ontology: Biological Process (GO: BP) analysis of macrophages after CD3ζ ICD signal transduction. **(b)** Flow cytometry analysis of PD-L1 and H-2Kd expression of macrophages after CAR-mediated signal transduction with or without NF-kB inhibitor (pyrrolidine dithiocarbamate, PDTC, 100 μM).

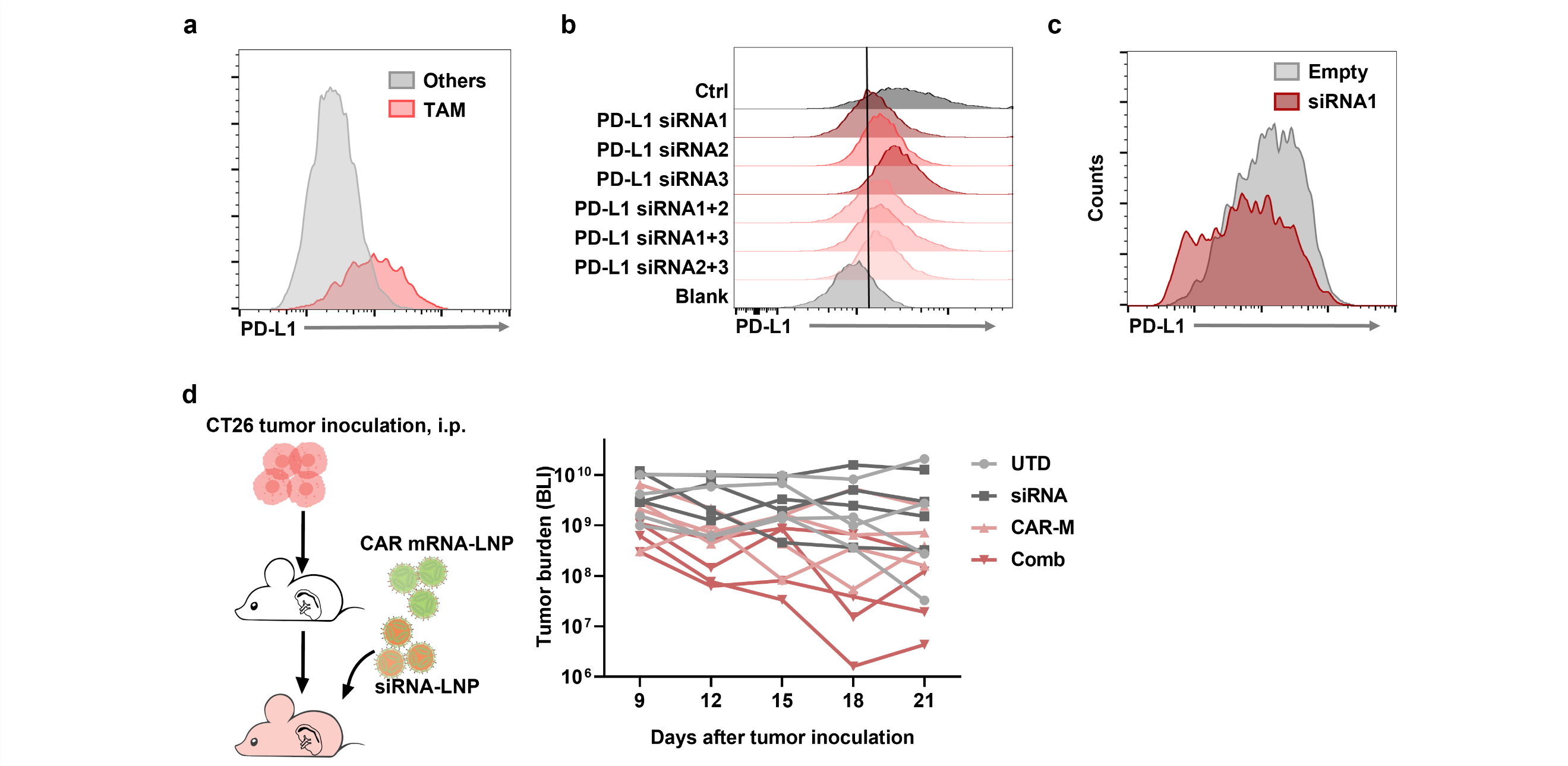

**Supplementary Fig. 18 | (a)** PD-L1 expression in TAMs and other cells from CT26-luc tumor. **(b)** Flow cytometry analysis of PD-L1 expression after PD-L1 siRNA (listed in Supplementary Table 3) knocking down *in vitro*. **(c)** Flow cytometry analysis of PD-L1 expression in TAM after PD-L1 siRNA knocking down *in vivo*. **(d)** Quantified BLI of the CT26-luc syngeneic mouse model with each treatment.

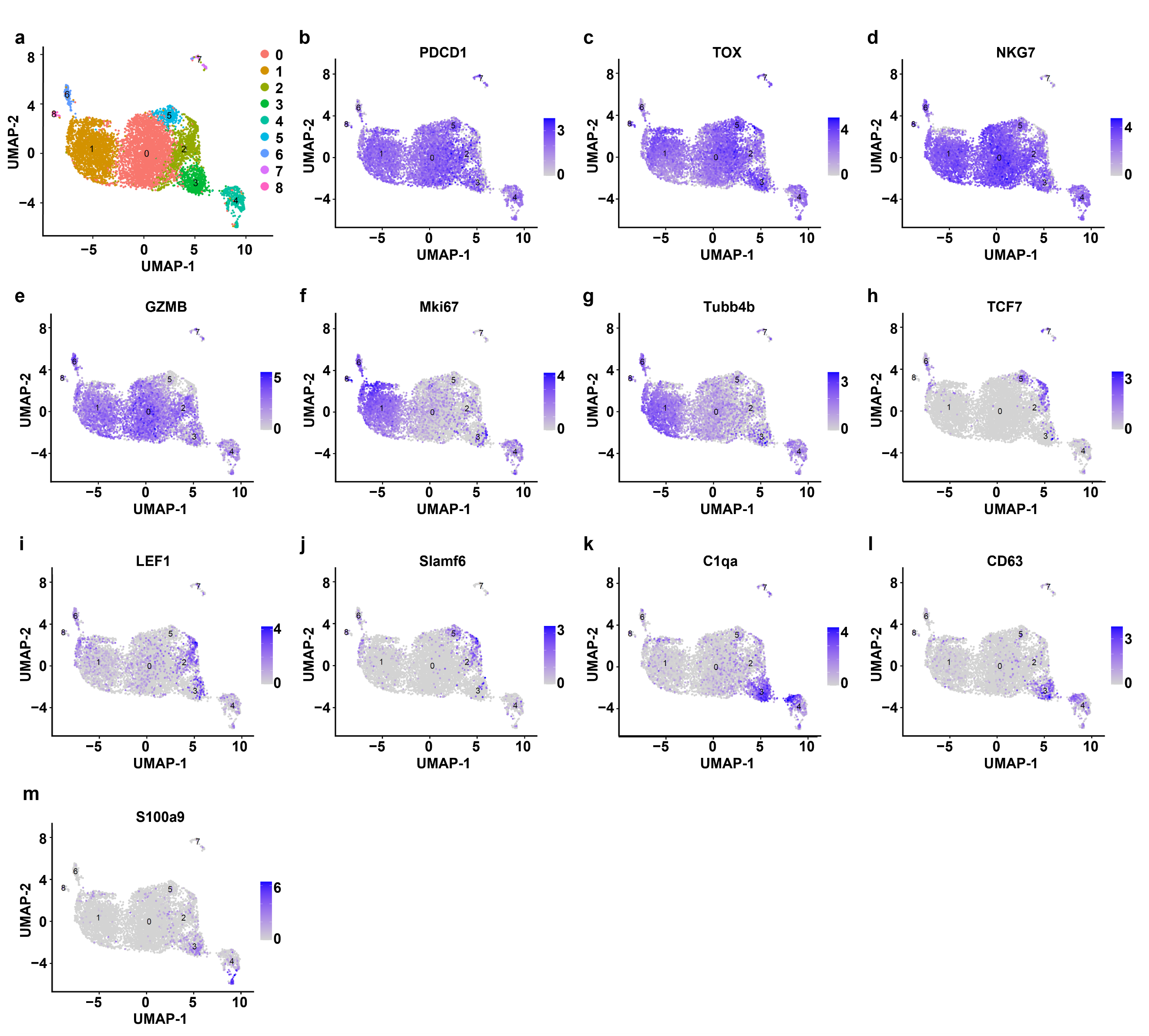

**Supplementary Fig. 19 | (a)** UMAP plot of scRNA-seq profiles from T cells within the tumor. UMAP plot of relative expression on the indicated genes (**(b)** PDCD1; **(c)** TOX; **(d)** NKG7; **(e)** GZMB; **(f)** Mki67; **(g)** Tubb4b; **(h)** TCF7; **(i)** LEF1; **(g)** Slamf6; **(k)** C1qa; **(l)** CD63; **(m)** S100a9).

**Supplementary Table 1.** CAR sequence studied in this work.

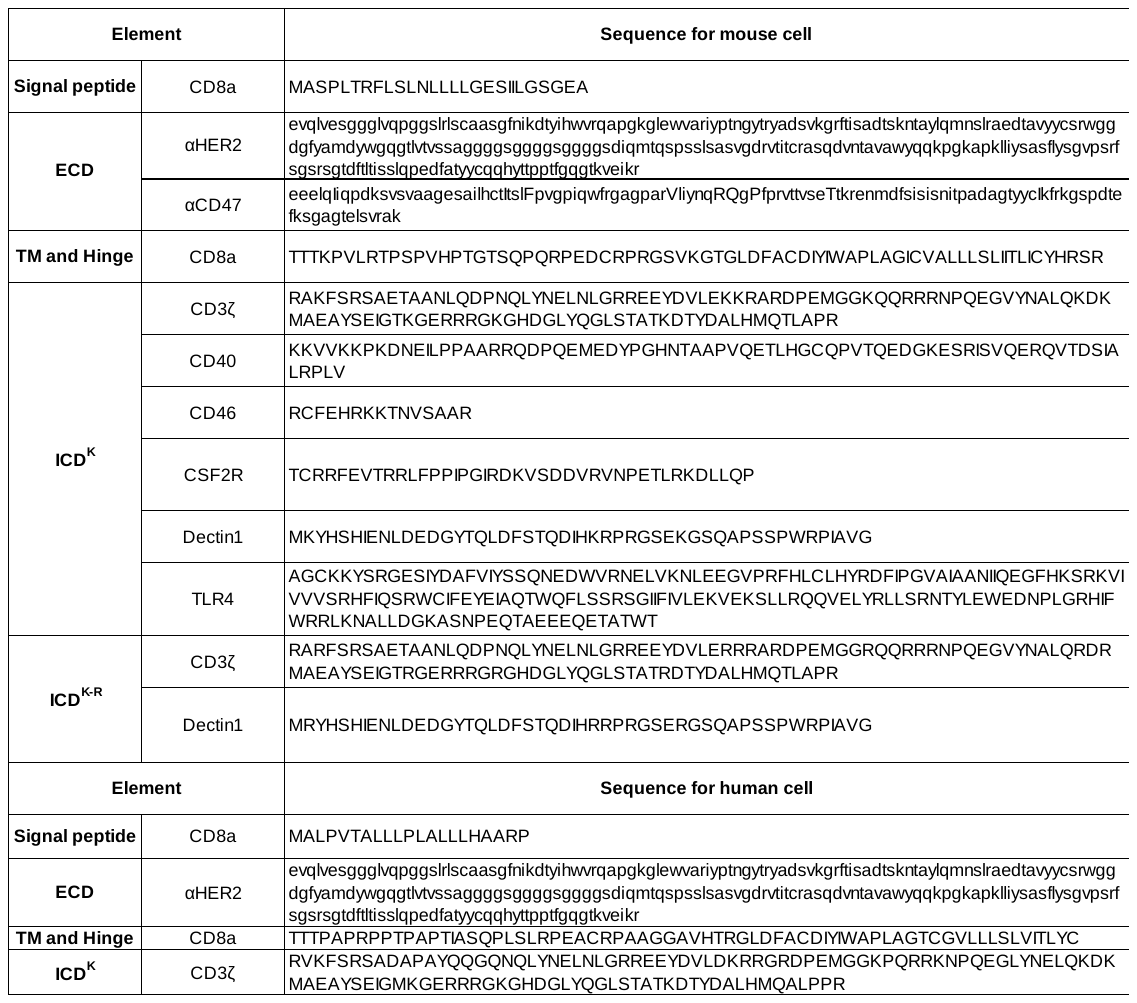

**Supplementary Table 2.** Blood count analysis data in healthy female Balb/c mice after the infusion of CAR-mRNA LNPs.

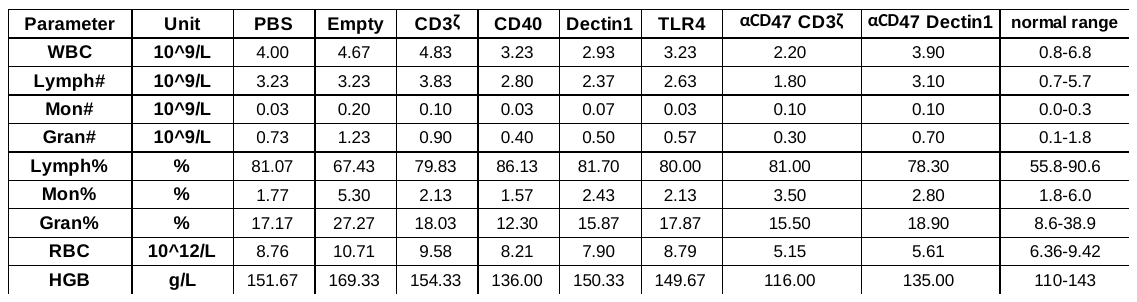

**Supplementary Table 3.** siRNA used in this study.

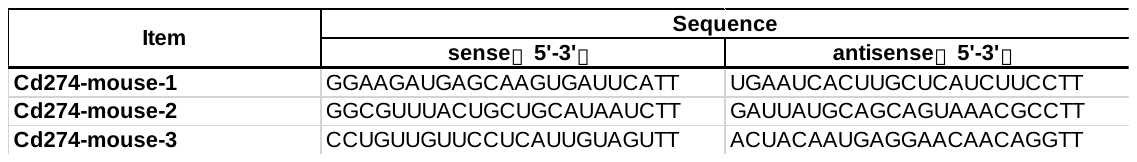

The siRNA1 used in the *in vivo* experiment was 2′-OMe-phosphorodithioate-modified.
